# The great divide: rhamnolipids mediate separation between *P. aeruginosa* and *S. aureus*

**DOI:** 10.1101/2023.06.23.546300

**Authors:** Jean-Louis Bru, Summer Kasallis, Rendell Chang, Quantum Zhuo, Jacqueline Nguyen, Phillip Pham, Elizabeth Warren, Katrine Whiteson, Nina Molin Høyland-Kroghsbo, Dominique H. Limoli, Albert Siryaporn

## Abstract

The coexistence of multiple bacterial species during infection can have significant impacts on pathogenesis. *Pseudomonas aeruginosa* and *Staphylococcus aureus* are opportunistic bacterial pathogens that can co-infect hosts and cause serious illness. The factors that dictate whether one species will outcompete the other or whether the two species can coexist are not fully understood. We investigated the role of surfactants in the interactions between these two species on a surface that enables *P. aeruginosa* to swarm. We found that *P. aeruginosa* swarms are repelled by colonies of clinical *S. aureus* isolates, creating physical separation between the two strains. This effect was abolished in mutants of *S. aureus* that were defective in the production of phenol-soluble modulins (PSMs), which form amyloid fibrils around wild-type colonies. We investigated the mechanism that establishes physical separation between the two species using the Imaging of Reflected Illuminated Structures (IRIS) method, which tracks the flow of the rhamnolipid surfactant layer produced by *P. aeruginosa*. We found that PSMs produced by *S. aureus* deflected the rhamnolipid surfactant layer flow, which in turn, altered the direction of *P. aeruginosa* swarms. These findings show that rhamnolipids mediate physical separation between *P. aeruginosa* and *S. aureus*, which enables these species to coexist in distinct microenvironments. Additionally, we found that a *Bacillus subtilis* surfactant and abiotic hydrophobic molecules repelled *P. aeruginosa* swarms through surfactant deflection. Our results suggest that surfactant interactions could have major impacts on bacteria-bacteria and bacteria-host relationships. In addition, our findings uncover a mechanism responsible for *P. aeruginosa* swarm development that does not rely on sensing but instead is guided largely by the flow of the surfactant layer and its boundaries.

## Introduction

*Pseudomonas aeruginosa* and *Staphylococcus aureus* are opportunistic pathogens that colonize the skin, eyes, and lungs, where they can become significant contributors to the development of a range of illnesses (Lyczak et al., 2000; Howden et al., 2023). While a single species can dominate over the other during the infection, co-infections have been reported to be associated with worse patient outcomes (Camus et al., 2022). In particular, the two species are commonly found together coinfecting the lungs of cystic fibrosis (CF) patients (Salsgiver et al., 2016). While the factors that impact the ability of these pathogens to compete or coexist are not fully understood, their molecular interactions could have an important role. Recent work has revealed that *P. aeruginosa* detects and responds to the presence of *S. aureus* in several ways. For example, the presence of *S. aureus* induces exploratory motility in *P. aeruginosa* microcolonies (Limoli et al., 2019) and *P. aeruginosa* detects *S. aureus* exoproducts including intermediate metabolites and molecules that modulate the iron starvation response (Zarrella and Khare, 2022).

Additional insight into how these species interact in host environments may be gained by considering their microenvironments. *P. aeruginosa* releases a number of molecules into its surroundings that improve its own fitness, including rhamnolipids (RLs), siderophores, and phenazines. The production of these molecules is regulated by the cell-cell communication process known as quorum sensing (QS), which enables groups of bacteria to coordinate collective behaviors by emitting and detecting QS molecules (Papenfort and Bassler, 2016). RLs have a significant impact on the *P. aeruginosa* microenvironment due to their surfactant properties, ubiquity, and multifunctional roles. RLs are glycolipids that consist of rhamnose and variable-length fatty acid moieties that are produced in high abundance (Jarvis and Johnson, 1949; Abdel-Mawgoud et al., 2010). They are amphipathic, containing both hydrophilic (rhamnose) and hydrophobic (fatty acid) groups, and function as surfactants, which decrease both the surface tension at air-liquid interfaces and the interfacial tension between two liquids. Due to their surfactant properties, RLs increase the solubility of hydrophobic molecules. This property improves the ability of *P. aeruginosa* to uptake otherwise poorly soluble molecules such as hydrocarbons, which can be used as nutrient sources (Beal and Betts, 2000; Noordman and Janssen, 2002; Abdel-Mawgoud et al., 2010). In addition, this property facilitates the solubilization of the *P. aeruginosa* QS molecule 2-heptyl-3-hydroxy-4-quinolone (PQS) (Collier et al., 2002; Calfee et al., 2005), which could affect cell-to-cell communication. RLs can directly impact *S. aureus* through the formation of micelles that transport toxins into the bacterium, altering its biofilm development and QS gene expression, and functioning as an antimicrobial (Haba et al., 2003; Benincasa et al., 2004; Gdaniec et al., 2022; Saadati et al., 2022). Serving as an important factor in pathogenesis, RLs are found in relatively high abundance in infection environments such as the lung, where they promote biofilm formation, and inhibit host immunity (Kownatzki et al., 1987; McClure and Schiller, 1992; Pamp and Tolker-Nielsen, 2007; Abdel-Mawgoud et al., 2010).

RLs have a critical role in the expansion of *P. aeruginosa* swarms, which are characterized by radially expanding dendritic patterns of densely packed cells that are referred to as tendrils (Caiazza et al., 2005). Cells within swarms are highly motile via flagellar activity, grow at a high density, and have high metabolic activity (reviewed in (Kearns, 2010)). Due to the rapid expansion of swarms across large distances, swarming has been widely described as a form of motility. RLs form a layer that expands outwards and precedes *P. aeruginosa* tendril formation (Siegmund and Wagner, 1991; Morris et al., 2011). The abundance of RLs and the spatial heterogeneity that are characteristic of swarms represent an ideal environment for the investigation of bacteria-bacteria interactions.

Recent work investigated the interaction between sub-populations of *P. aeruginosa* within swarms, finding that sub-populations stressed by phage infection or antibiotics reorganize *P. aeruginosa* tendrils (Bru et al., 2019). This reorganization creates a spatial separation between stressed and untreated populations, enabling “healthy” *P. aeruginosa* populations to physically avoid agents that induce stress. This collective stress response occurs through altered production of RLs and the QS-associated molecules PQS and 2-heptyl-4-quinolone (HHQ) (Fig. 1A) (Bru et al., 2019). Untreated populations produce RLs, which facilitate swarming. Phage or antibiotic-stressed populations decrease production of RLs and increase the production of PQS, which signals the redirection of healthy tendrils away from the area of stress (Bru et al., 2019). The same process could also facilitate physical separation of *P. aeruginosa* from predators and competitors. In particular, this separation mechanism could facilitate coexistence between *P. aeruginosa* and other bacterial species. Many questions remain regarding how tendrils are steered away from the stressed populations. For example, *P. aeruginosa* can sense PQS produced by the stressed populations through the QS regulator PqsR (Wade et al., 2005). However, the mechanism by which the swarm alters its tendril motion in response to the detection of PQS is unknown. Additionally, it is not understood how prevalent the collective stress response is among strains of *P. aeruginosa* or other bacterial species including *S. aureus*.

**Figure 1.**
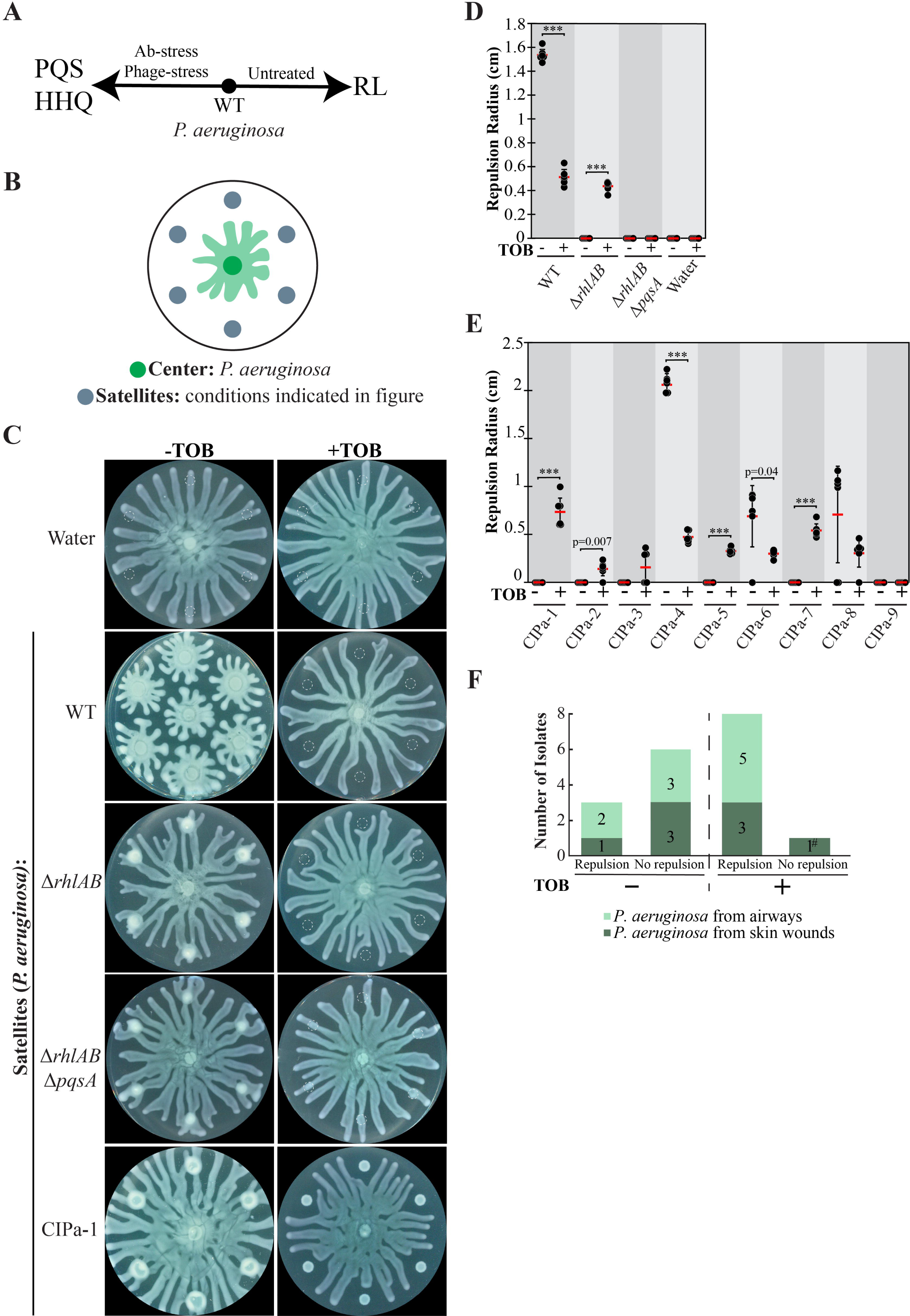
The effect of tobramycin on swarm repulsion by P. aeruginosa WT, mutants, and clinical isolates. (A) Schematic depicting relationship between the QS molecule 2-heptyl-3-hydroxy-4-quinolone (PQS) and rhamnolipid (RL) production in *P. aeruginosa* on swarming plates. Untreated *P. aeruginosa* produces more RLs whereas antibiotic- or phage-stressed cells produce more PQS. (B) Schematic depicting swarm interaction assays in which *P. aeruginosa* is inoculated at the center and forms swarm tendrils that move outwards. (C) Swarm interaction assays in which wild-type (WT) *P. aeruginosa* was spotted at the center and test strains were spotted at satellite positions. Tobramycin (TOB) treatment was performed by mixing TOB with bacteria to a final concentration of 0.5 mg/mL and spotting 6 μL of the mixture onto the swarm plate. Images were acquired 18-20 hours following inoculation. Dashed lines indicate the boundaries of the initial inoculum. Quantification of tendril repulsion by (D) *P. aeruginosa* PA14 WT and mutant strains and (E) *P. aeruginosa* clinical isolates from airways or skin wounds. Black dots represent the repulsion radius (cm) of individual satellites and red bars indicate the average. Error bars indicate standard deviations. T-tests were performed as two-tailed distributions with unequal variance, *** denotes p < 0.001. (F) Plot summarizing the number of clinical isolates that either repelled or did not repel swarm tendrils in swarm interaction assays. Isolates were considered repulsive if at least three positions repelled the center swarm. Isolates were TOB-treated or untreated. The hash mark (#) denotes the TOB-resistant isolate. Swarm assay images for the clinical isolates are shown in Fig. S1A in the Supplementary Materials.

*S. aureus* modifies its microenvironment through the production of phenol-soluble modulins (PSMs), which are amphipathic peptides that have surfactant properties, antimicrobial activity, and inhibit host innate immunity (Kaito and Sekimizu, 2007; Cogen et al., 2010; Tsompanidou et al., 2011; Peschel and Otto, 2013). Interestingly, RLs produced by *P. aeruginosa* share these same properties. Multiple PSMs have been identified, including PSMα1 to PSMα4 (∼20 residues), PSMβ1 and PSMβ2 (∼40 residues), and PSMγ, also referred to as δ-toxin (Wang et al., 2007; Peschel and Otto, 2013). δ-toxins are similar to PSMα and are encoded by the *hld* gene. PSMs can aggregate, which results in the formation of amyloid fibrils that have multiple functions, including the fortification of the matrix of *S. aureus* biofilms and increasing the cytotoxicity of PSMs (Schwartz et al., 2012; Tayeb-Fligelman et al., 2017; Zheng et al., 2018; Zaman and Andreasen, 2020; Kreutzberger et al., 2022). PSMs facilitate the expansion of *S. aureus* on surfaces through spreading and comet formation, a mechanism similar to the active motility mechanism known as gliding (Tsompanidou et al., 2013; Pollitt et al., 2015). The functional roles of PSMs and RLs and their impact on their microenvironments suggest that they could have a significant role in *P. aeruginosa* - *S. aureus* interactions.

Here, we investigate how *P. aeruginosa* swarm tendrils interact with *S. aureus* clinical isolates. We find that clinical strains of *P. aeruginosa* that are stressed by the antibiotic tobramycin repel swarm tendrils, which demonstrates the ubiquity of the collective stress response. Untreated *S. aureus* can repel *P. aeruginosa* swarm tendrils, an effect which is abolished in *S. aureus* strains that are defective in the production of PSMs. We use IRIS imaging to track the movement of surfactant, which is dependent on rhamnolipid synthesis. We find that the presence of *S. aureus* alters the flow of surfactant in *P. aeruginosa* swarms, creating a zone of surfactant exclusion which results in tendril repulsion. This effect is significantly reduced in *S. aureus* PSM mutants. Our results show that the surfactant layer, which is composed of RLs, mediates the repulsion of *P. aeruginosa* swarms away from *S. aureus* and multiple molecules including PQS and the surfactin produced by *B. subtilis*.

## Results

### Collective stress response in P. aeruginosa clinical isolates

The collective stress response was previously reported in a limited number of *P. aeruginosa* strains (Bru et al., 2019). We investigated the prevalence of the collective stress response in 9 clinical isolates of *P. aeruginosa* using swarm interaction assays in order to determine if this response could be of clinical relevance. In these assays, *P. aeruginosa* strain PA14 was inoculated as a single spot at the center of a swarm plate containing swarm medium and test strains were inoculated at six satellite positions surrounding the center spot (Fig. 1B). As the center swarm expands radially, the swarm tendrils approach and interact with the strain at the satellite positions (Fig. 1B). This swarm interaction assay enables replication of a single condition or the testing of multiple conditions on the same swarm plate. When *P. aeruginosa* strain PA14 is spotted at satellite positions, it repels the center swarm due to the production of RLs (Bru et al., 2019), an effect that is abolished in Δ*rhlAB* strains, which do not produce RLs (Fig. 1C-D). To elicit the collective stress response, antibiotics were supplied at satellite positions at above-MIC concentrations that slow growth but do not entirely inhibit it due to the diffusion of antibiotics away from the initial spot. Previous work showed that a number of antibiotics including gentamicin, kanamycin, and fosfomycin, elicited the collective stress response, causing antibiotic-stressed strains at satellite positions to repel untreated swarms (Bru et al., 2019).

Clinical isolates of *P. aeruginosa* were isolated from airway or skin infections. The isolates exhibited varying degrees of antibiotic resistance, but were largely sensitive to tobramycin (TOB), as demonstrated by the inhibition of growth or swarming in TOB-treated satellite colonies (Fig. 1C, 1E-F and Fig. S1A in Supplementary Materials). TOB targets the bacterial 30S ribosomal subunit, blocks tRNA translocation (Ying et al., 2019), and induces the production of PQS in *P. aeruginosa* (Morales-Soto et al., 2018; Rieger et al., 2020). TOB-treated WT and Δ*rhlAB* strains at satellite positions were significantly inhibited for growth but repelled the untreated center swarm (Fig. 1C-D). This result is consistent with TOB triggering the PQS-mediated collective stress response. We note that PQS alone is sufficient to repel the center swarm ((Bru et al., 2019) and Fig. S1B in Supplementary Materials). TOB-treatment of *P. aeruginosa* Δ*rhlAB* Δ*pqsA*, which lacks the ability to produce both RLs and PQS, failed to elicit tendril repulsion (Fig. 1C-D), further supporting the dependence of the TOB-induced repulsive response on PQS. Center swarm tendrils collided with satellite positions containing only TOB, indicating that the repulsion is not due to the presence of TOB alone (Fig. 1C-D). Together, the results demonstrate that TOB treatment of *P. aeruginosa* stimulates the collective stress response, which repels swarms through the production of PQS. These results further bolster the model that *P. aeruginosa* swarms produce RLs in non-stressed conditions but increase PQS production under stress from antibiotics (Fig. 1A).

We assessed the TOB-induced collective stress response in the clinical isolates. Colonies were considered to repel the center swarm if the tendrils did not come into contact with three or more of the satellite inoculum positions. In the absence of TOB, three of the clinical isolates (CIPa-4, CIPa-6, CIPa-8) repelled the center swarm (Fig. 1E-F and Fig. S1A in Supplementary Materials). Treatment with TOB significantly reduced *P. aeruginosa* growth and caused repulsion of the swarm tendrils in 9 out of the 10 clinical isolates (Fig. 1E-F and Fig. S1A in Supplementary Materials). The only strain that did not repel when treated with TOB was CIPa-9, which appeared to be TOB-resistant. The significant increase in the number of strains that repelled swarms due to TOB treatment (Fig. 1F) is consistent with activation of a collective stress response and suggests that PQS is produced in response to antibiotic stress in the TOB-sensitive clinical isolates tested. Together, these results demonstrate that the collective stress response is present in *P. aeruginosa* clinical isolates that cause disease in human airways and skin wounds.

### S. aureus clinical isolates repel P. aeruginosa swarms

The collective stress response in *P. aeruginosa* could promote its survival in natural environments by separating the swarm population from areas that contain phage or antibiotics. We reasoned that *P. aeruginosa* could use a similar strategy to isolate itself from other bacterial species. We assessed the interaction between *P. aeruginosa* and *S. aureus* by inoculating *P. aeruginosa* at the swarm plate center and *S. aureus* at satellite positions (Fig. 2A). Under these conditions, *P. aeruginosa* developed tendrils that moved outwards whereas *S. aureus* grew as colonies that did not expand spatially. We tested the interaction of the methicillin-resistant *S. aureus* clinical isolates USA300, USA300 JE2, and USA300 (LAC) and found that these strains strongly repelled *P. aeruginosa* tendrils (Fig. 2B-C), similar to antibiotic-stressed *P. aeruginosa*. Treatment of the *S. aureus* strains with TOB reduced colony growth, similar to the effect on *P. aeruginosa* strains (Fig. 1C, 2B). However, this treatment of *S. aureus* also suppressed the repulsion of *P. aeruginosa* tendrils and enabled the tendrils to infiltrate the *S. aureus* colonies (Fig. 2B-C). *P. aeruginosa* repulsion by untreated *S. aureus* and a lack of repulsion by TOB-treated *S. aureus* was largely observed in clinical isolates of *S. aureus*, which were isolated from airways and skin wounds. In 9 out of the 10 clinical isolates, untreated *S. aureus* strains repelled *P. aeruginosa* tendrils, whereas TOB-treated strains suppressed tendril repulsion (Fig. 2D-E and Fig. S2 in Supplementary Materials).

**Figure 2.**
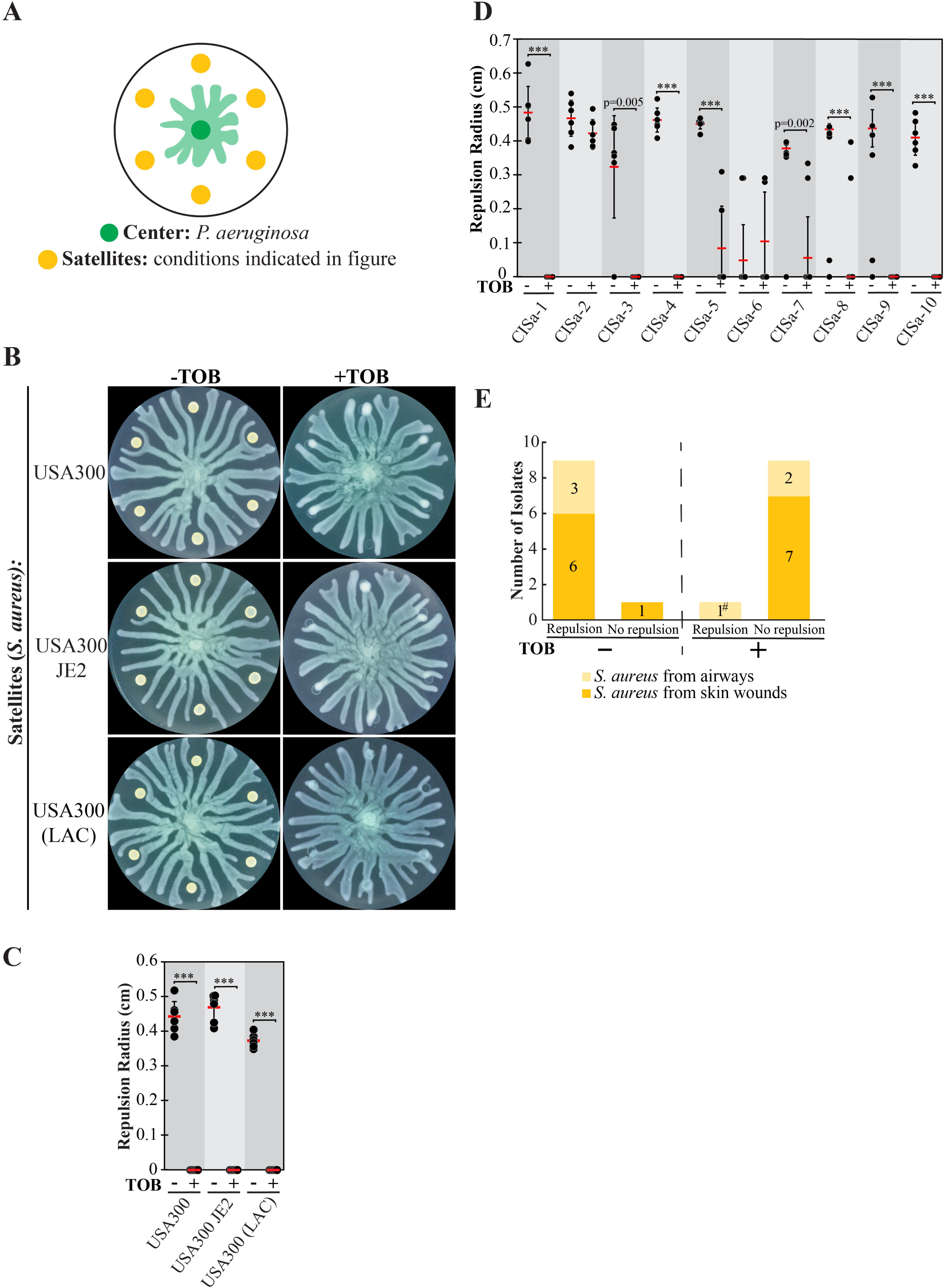
The effect of tobramycin on swarm repulsion by *S. aureus* WT and clinical isolates. (A) Schematic indicating swarm interaction assay in which *P. aeruginosa* and *S. aureus* are inoculated at the center and satellite positions respectively, and *P. aeruginosa* swarm tendrils move outward towards the satellite positions. (B) Swarm interaction assays in which WT *P. aeruginosa* and WT *S. aureus* strains were spotted at satellite positions. TOB treatment was performed by mixing TOB with bacteria to a final concentration of 0.5 mg/mL and spotting 6 μL of the mixture onto the swarm plate. Plates were imaged 18-20 hours following inoculation. Quantification of repulsion by (C) *S. aureus* USA300 strains and (D) *S. aureus* clinical isolates from airways or skin wounds. Black dots represent the repulsion radius (cm) of individual satellites and red bars indicate the average. Error bars indicate standard deviations. (E) Plot summarizing the number of clinical isolates that either repelled or did not repel *P. aeruginosa* swarm tendrils in swarm interaction assays with or without TOB treatment. Isolates were considered repulsive if at least three positions repelled the center swarm. The hash mark (#) denotes the TOB-resistant isolate. T-tests were performed as two-tailed distributions with unequal variance, *** denotes p < 0.001. Swarm assay images for the clinical isolates are shown in Fig. S2 in the Supplementary Materials.

Whereas antibiotic treatment of *P. aeruginosa* caused tendril repulsion, this treatment of *S. aureus* suppressed tendril repulsion. Thus, *P. aeruginosa* is repelled by antibiotics stress of its own species, but not by the antibiotic stress of its competitor *S. aureus*. This suggests that the collective stress response is beneficial for only its own species. These findings raise the question of how *S. aureus* repels *P. aeruginosa* tendrils. While tendrils are repelled by antibiotic-stressed *P. aeruginosa* via the production of the QS molecule PQS, *S. aureus* does not contain the genes necessary for PQS synthesis. We hypothesized that *S. aureus* produces another QS-associated molecule that is sensed by *P. aeruginosa,* which ultimately alters tendril direction.

### PSM fibrils facilitate repulsion of P. aeruginosa swarms

We investigated the potential role of the QS-regulated phenol soluble modulins (PSMs) produced by *S. aureus* in the repulsion of *P. aeruginosa* tendrils. We assessed the extent of repulsion by the *S. aureus* Δ*psmα* mutant, which still produces the δ-toxin and PSMβ. Whereas WT *S. aureus* strongly repelled *P. aeruginosa* swarms (Fig. 2B-C), the Δ*psmα* mutant was deficient in repulsion, with *P. aeruginosa* tendrils infiltrating these colonies (Fig. 3A). The Δ*psmα* Δ*hld* mutant, which does not produce any PSMα peptides or the δ-toxin, was also deficient in repelling *P. aeruginosa* swarms (Fig. 3A). This was observed in both the USA300 and USA300 (LAC) backgrounds. These findings suggest that PSMα peptides significantly contribute to the repulsion of *P. aeruginosa* swarms.

**Figure 3.**
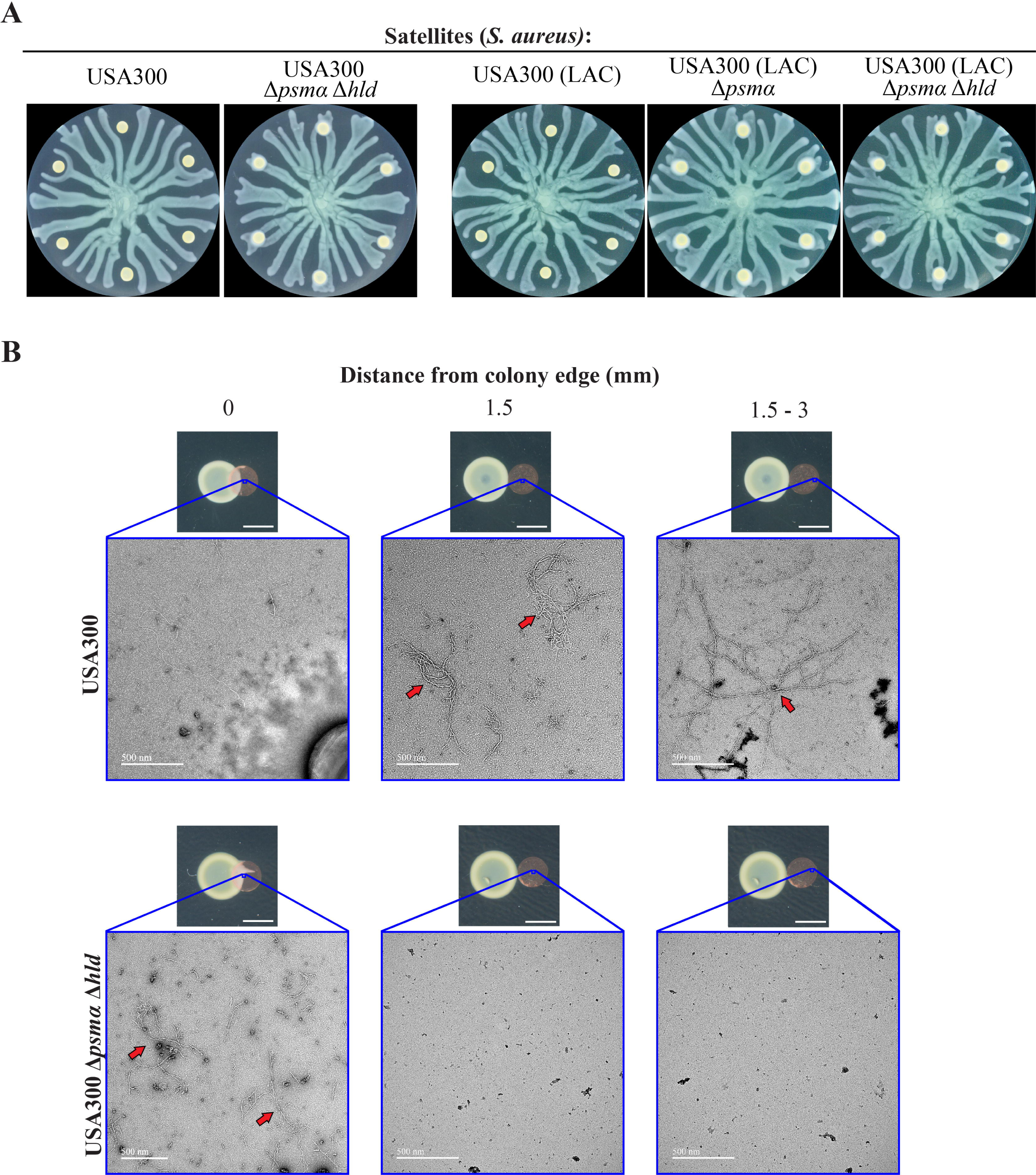
Dependence of *P. aeruginosa* tendril repulsion on phenol-soluble modulins (PSMs). (A) Swarm interaction assays in which WT *P. aeruginosa* was spotted at the center and WT *S. aureus* (USA300 and USA300 (LAC)) or the *S. aureus* Δ*psmα* or Δ*psmα* Δ*hld* mutants were spotted at the satellite positions. The images for WT *S. aureus* are the same as those in Fig. 2B and are shown here for reference. (B) Transmission electron microscopy (TEM) images of copper grids that were placed at or adjacent to inoculums of WT *S. aureus* or the Δ*psmα* Δ*hld* mutant on swarm plates, incubated 18 - 20 hours, and harvested. The relative positions of the TEM grid and *S. aureus* colony are shown (small images). TEM images (large images) were acquired from the approximate location indicated on the TEM grids. Fibrils (red arrows) were observed at all positions on the grids in the vicinity of the WT *S. aureus* colony. No fibrils were observed at positions that were 1.5 mm or greater from the edge of the *S. aureus* Δ*psmα* Δ*hld* colony. Scale bars indicate 3 mm and 500 nm in the small and large images, respectively.

*S. aureus* PSMs can form amyloid fibrils that support biofilm structures (Tayeb-Fligelman et al., 2017; Zheng et al., 2018; Zaman and Andreasen, 2020; Kreutzberger et al., 2022). We considered that PSM fibrils could mediate *P. aeruginosa* repulsion. First, we investigated the formation of fibrils by PSMs on the surface of the swarm plate surface using transmission electron microscopy (TEM). Carbon-coated copper TEM grids were placed at or adjacent to *S. aureus* inoculation sites at satellite positions and were incubated for the same period of time as the swarming assays to allow for PSM production and spreading. In WT *S. aureus*, we observed fibrous structures that were consistent with amyloid fibrils. The fibrils were present at distances up to 2.5 mm from the colony edge (Fig. 3B and Fig. S3A in Supplementary Materials). In contrast, no fibrils were observed in the Δ*psmα* Δ*hld* mutant beyond the colony edge (Fig. 3B and Fig. S3B in Supplementary Materials). These observations suggest PSMs form a layer of amyloid fibrils that surrounds WT *S. aureus* colonies that repels *P. aeruginosa* tendrils.

### Deflection of the surfactant layer causes tendril repulsion

We probed deeper into the mechanism underlying the repulsion of tendrils by PSM amyloid fibrils. We considered two models which could explain tendril repulsion by PSMs: (1) biological sensing of the PSMs and (2) a physicochemical mechanism that alters the development of tendrils. In the first model, *P. aeruginosa* could sense PSM fibrils and alter its motility away from them. For example, the *P. aeruginosa* pilus chemoreceptor PilJ is involved in sensing *S. aureus* and it is possible that it could detect PSMs (Yarrington et al., 2022). In the second model the repulsion of tendrils does not require detection by *P. aeruginosa*, but rather is due to physicochemical interactions between the amphipathic PSMs and the swarm. In this model, PSMs form a fibril boundary around *S. aureus* that repels approaching tendrils by altering the spatial distribution of the RLs near the *P. aeruginosa* tendrils.

To differentiate between these two models and better understand tendril development, we performed a series of experiments using abiotic molecules. We reasoned that tendrils would not be repelled by abiotic molecules if biological sensing of molecules is the predominant mechanism. We selected non-polar molecules with low volatilities to ensure that they remained on the swarming plate throughout the duration of the assays. Long-chain carbon molecules fit these criteria, including: oleic acid and linoleic acid (fatty acids), glyceryl trioleate and glyceryl trilinoleate (triglyceride forms of the fatty acids), Triton X-100 and Tween-20 (surfactants), and the liquid lubricant polydimethylsiloxane (PDMS) (Fig. 4A).

**Figure 4.**
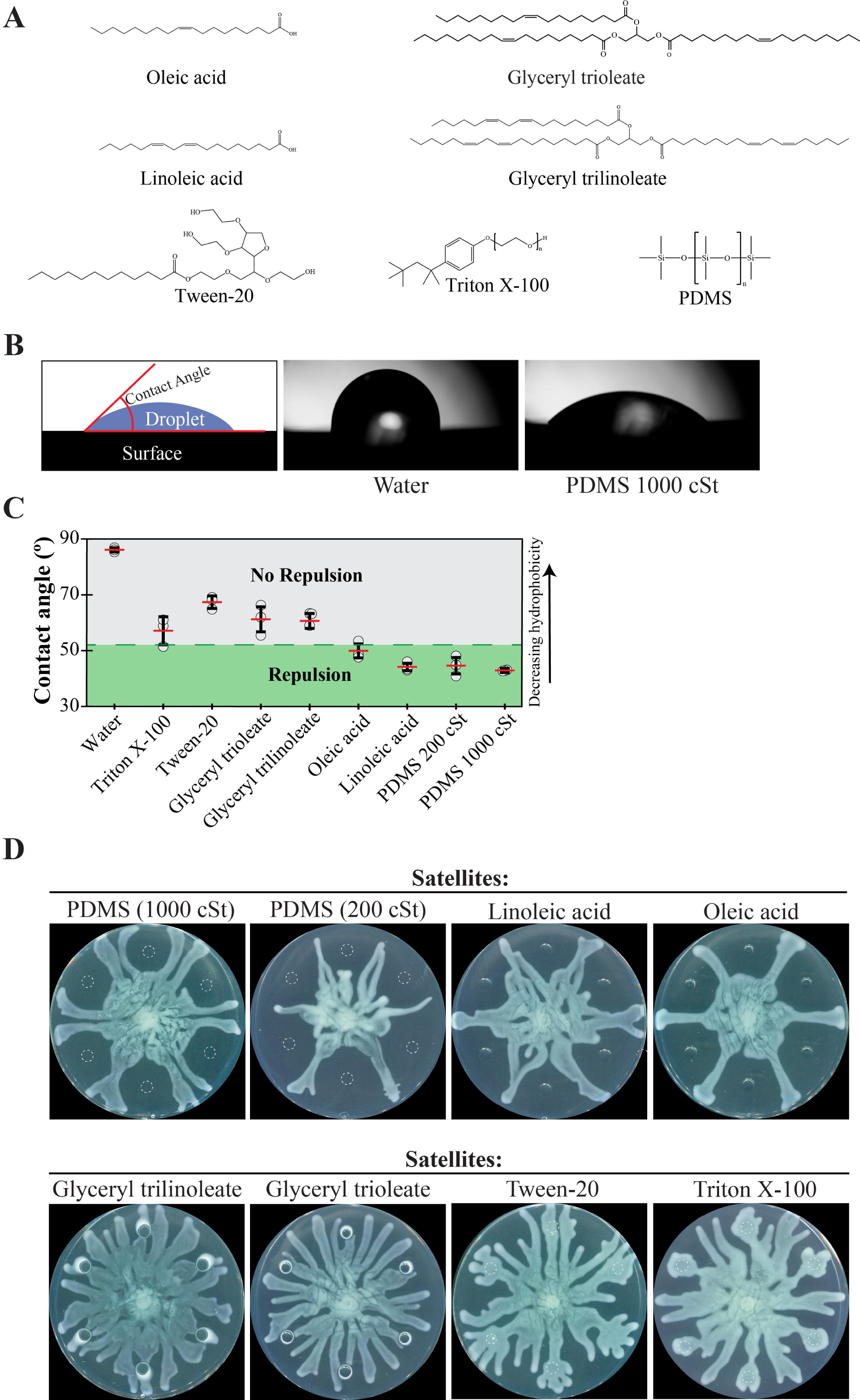
Repulsion of *P. aeruginosa* swarms by hydrophobic molecules. (A) Chemical structures of hydrophobic molecules that were tested for repulsion in swarm interaction assays. (B) Schematic indicating how contact angles were measured using images from a contact angle goniometer. Images of droplets on an oleophobic surface are shown for water and 1000 cSt PDMS, which gave the highest and lowest contact angles, respectively. (C) Contact angle measurements for each of the molecules. Repulsion or lack of repulsion was assessed using the data in Fig. 4D and 1C. Points indicate the average measurement of the left and right sides of each droplet. Red lines indicate the average (n=6) and error bars indicate standard deviation. Representative droplet images can be found in Fig. S4 in the Supplementary Materials. (D) Swarm interaction assays in which WT *P. aeruginosa* and test molecules were spotted at the center or satellite positions, respectively. Images were acquired 15 hours following inoculation. Triton X-100 and Tween-20 were used at concentrations of 0.2% and 2%, respectively, due to their inhibitory effect on *P. aeruginosa* growth at higher concentrations.

To determine the relative hydrophobicity of each molecule, we measured its contact angle on an oleophobic surface using a contact angle goniometer. This device measures the angle that a droplet makes with the surface and gives a quantitative measure of hydrophobicity (Fig. 4B). Oleophobic surfaces maintain droplet forms for both hydrophobic and hydrophilic molecules, which allows for contact angle measurements for a wide range of hydrophobicities using a single surface. PDMS had the smallest contact angle, indicating the greatest hydrophobicity of the molecules tested (Fig. 4C). In order of decreasing hydrophobicity (increasing contact angle) were PDMS, the fatty acids, triglycerides, and the surfactant Tween-20 (Fig. 4C and Fig. S4 in Supplementary Materials). Water was included as a reference and had the greatest contact angle. We spotted 6 µL of each molecule at satellite positions on swarm plates at the beginning of swarm interaction assays. If our hypothesis that tendril repulsion does not require sensing is correct, tendrils would be repelled by the hydrophobic molecules. In support of the hypothesis, PDMS and the fatty acids repelled tendrils (Fig. 4D). Triglycerides and surfactants, which are less hydrophobic, did not repel tendrils (Fig. 4D). In addition, we tested the effect of viscosity on tendril repulsion through the use of PDMS at two different viscosities, 200 cSt and 1000 cSt. For reference, water has a viscosity of 1 cSt at 20°C. The lower viscosity (200 cSt) PDMS produced greater repulsion than the higher viscosity PDMS (1000 cSt) (Fig. 4D and S4B in Supplementary Materials), suggesting that a lower viscosity molecule of the same hydrophobicity produces greater repulsion. Together, these results show that hydrophobic molecules can repel swarm tendrils.

We next hypothesized that hydrophobic molecules repel tendrils by altering the spatial organization of surfactant that is produced by the swarm. The surfactant contains RLs and precursor 3-(3-hydroxyalkanoyloxy) alkanoic acids (HAAs), which facilitate tendril development (Caiazza et al., 2005; Soberón-Chávez et al., 2005; Abdel-Mawgoud et al., 2010). Tracking the spatial distribution of surfactants has been a significant challenge because they are optically transparent. In literature, surfactants from swarms have been visualized using methylene blue and Nile red dyes (Siegmund and Wagner, 1991; Morris et al., 2011) but these dyes can alter surfactant and swarm dynamics. To better measure the dynamics of surfactants and swarm tendrils without perturbing them, we used an imaging method we recently developed, Imaging of Reflected Illuminated Structures (IRIS) (Kasallis et al., 2023). IRIS illuminates swarm surfaces by projecting a structured image onto the surface and capturing the reflection (Fig. 5A). The projected structured image is a checkerboard pattern, which increases the contrast of surface features, most notably the liquid-air interface at the edge of the swarm. Conventional ambient lighting or reflection of a non-structured image captures the boundaries of tendrils but the surfactant is not discernible (Fig. 5B). In contrast, IRIS imaging reveals high resolution details about the swarm. Notably the tendrils rest on a distinct optically transparent fluidic layer (Fig. 5B). We used edge detection to further demarcate the tendrils and fluidic layers. The fluidic layer is not present in the Δ*rhlAB* mutant, which does not produce the HAAs or RLs (Fig. S5A in Supplementary Materials). This observation suggests that the fluidic layer is composed of HAAs and RLs, which together form a surfactant layer. We performed differential analysis of timelapse IRIS images to identify the components of the swarm that were dynamic while suppressing static components (Fig. 5B). This analysis showed previously that the surfactant layer expands in lockstep with the tendril edges, which suggests that movement of the surfactant layer and tendrils are coupled (Kasallis et al., 2023). Building upon this finding, we considered the possibility that hydrophobic molecules could alter the flow of the surfactant layer, which in turn could alter the direction of tendrils.

**Figure 5.**
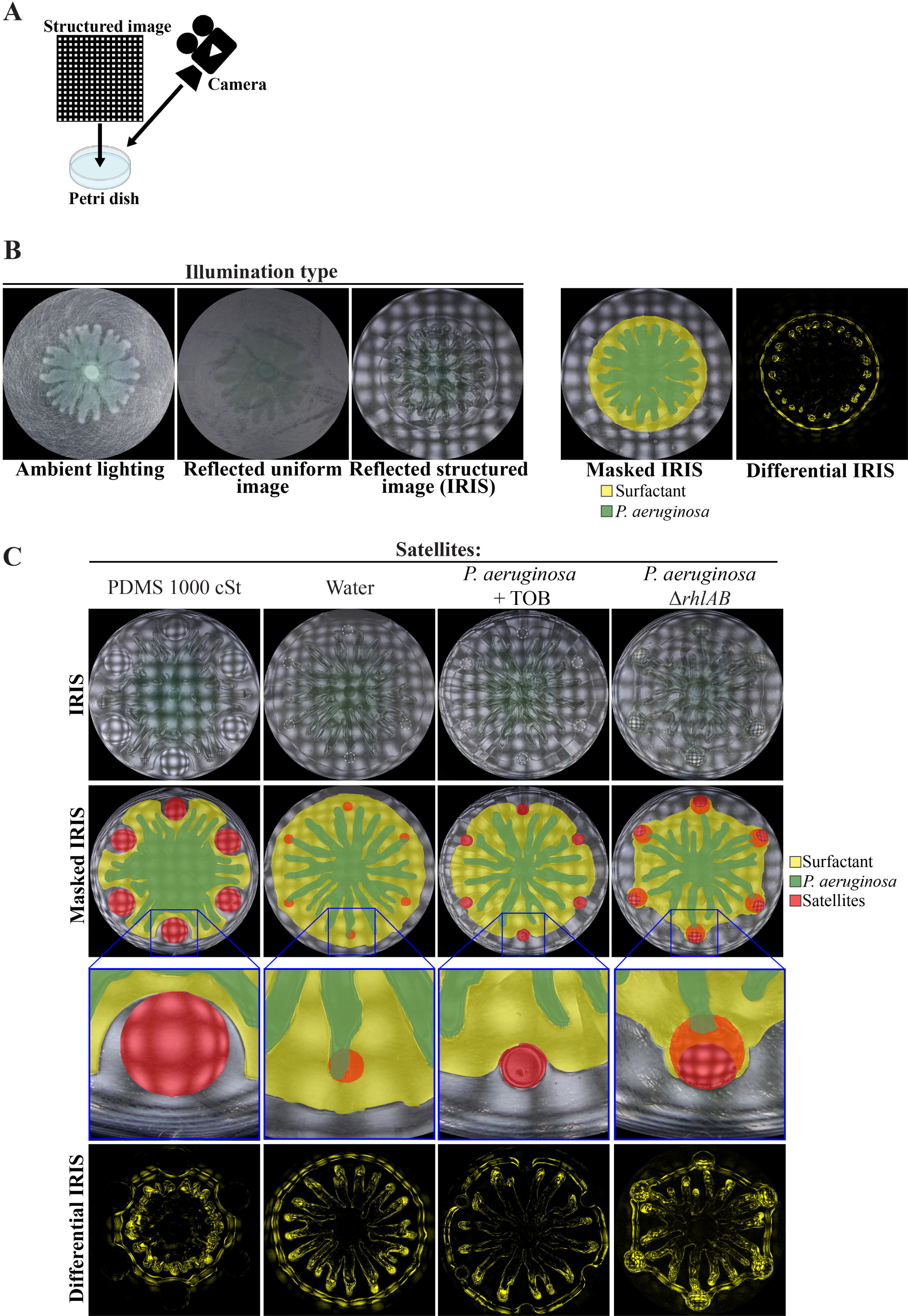
Role of the *P. aeruginosa* surfactant layer in swarm repulsion. (A) Schematic of the Imaging using Reflected Illuminated Structures (IRIS) setup. (B) Swarm assay in which WT *P. aeruginosa* was spotted at the center and imaged using ambient lighting, a reflected uniform image, or a reflected structured image at 11 hours following inoculation. The masked IRIS image indicates the surfactant layer (yellow) and *P. aeruginosa* (green) boundaries. The differential IRIS image indicates components of the surfactant layer and *P. aeruginosa* that are dynamic. (C) IRIS, masked IRIS, and differential IRIS images of swarm interaction assays in which WT *P. aeruginosa* was spotted at the center and either test molecules (1000 cSt PDMS or water) or strains (TOB-treated *P. aeruginosa* or *P. aeruginosa* Δ*rhlAB*) were spotted at satellite positions. Images were acquired 12-14 hours following inoculation. Masked IRIS images indicate the surfactant layer (yellow), *P. aeruginosa* (green), and test molecules or strains at the satellite positions (red). In *ΔrhlAB*, the surfactant layer overlaps the Δ*rhlAB* satellite colonies. Dashed lines at satellite positions show the boundary of the initial spot. Magnified IRIS images that are not masked can be found in Fig. S5B in the Supplementary Materials.

We imaged the dynamics of the *P. aeruginosa* surfactant layer and its interaction with the hydrophobic molecules using IRIS timelapse imaging. PDMS, which was the most hydrophobic molecule that we tested (Fig. 4C), strongly deflected the surfactant layer, causing the surfactant to flow clearly around the PDMS (Fig. 5C, and Fig. S5B and Movie S1 in Supplementary Materials). As expected, the surfactant layer and tendrils were not deflected by water (Fig. 5C and Fig. S5B in Supplementary Materials). Concomitant with the deflection of the surfactant, the tendrils moved around the PDMS. A notable feature of the dynamics is that the tendrils were constrained to move within the boundary of the surfactant layer. These observations raise the possibility that the surfactant layer sets the boundaries on which the tendrils can move, which constrains and alters the tendril movements. It is unlikely that *P. aeruginosa* could alter the direction of the surfactant flow to cause the surfactant to move around the PDMS. A more plausible explanation is that the hydrophobicity of the PDMS deflects the surfactant layer, causing the surfactant to move around the PDMS. This interpretation suggests that the physicochemical interaction between the surfactant layer and PDMS is responsible for tendril repulsion by PDMS, though the interpretation does not entirely rule out a role for sensing of PDMS by *P. aeruginosa*.

We hypothesized that biological molecules, including PQS produced by stressed *P. aeruginosa* populations (Fig. 1C-D and Fig. S1B in Supplementary Materials), can repel tendrils through surfactant deflection. In support of this hypothesis, IRIS imaging indicated that surfactant from the *P. aeruginosa* swarm was indeed deflected by TOB-treated *P. aeruginosa* satellite populations (Fig. 5C). The movement of swarm tendrils away from TOB-treated populations was concomitant with the deflection of the surfactant (Movie S2 in Supplementary Materials). These data suggest that tendril repulsion by antibiotic-stressed *P. aeruginosa* is caused by deflection of the surfactant layer, which in turn, alters the movement of swarm tendrils around the stressed *P. aeruginosa* population. We verified that *P. aeruginosa* alone does not repel tendrils or deflect the surfactant layer by using the Δ*rhlAB* strain, which does not produce PQS without antibiotic stress and is deficient in RL production (Bru et al., 2019). Surfactant produced by the WT swarm first merged with Δ*rhlAB* colonies at the satellite positions, followed by the advancement of swarm tendrils into the satellite boundaries (Fig. 5C and Movie S3 in Supplementary materials).

The effects on the surfactant layer are prominent in differential IRIS images, which show that the surfactant boundary curves inward into a concave shape as it approaches PDMS or TOB-treated *P. aeruginosa* (Fig. 5C). In contrast, no concave features are observed in the surfactant layer as it approaches the Δ*rhlAB* mutant. In addition, the surfactant appears to be attracted to the Δ*rhlAB* colony, causing the surfactant layer to develop into a hexagonal pattern (Fig. 5C and Movie S3 in Supplementary Materials). We note that surfactant flow from the WT swarm is observed within the boundary of the Δ*rhlAB* satellite colony, as detected by the differential IRIS images. This flow is not observed in the TOB-treated WT colonies (Movie S2 in Supplementary Materials) and is consistent with merging of the surfactant layer with the Δ*rhlAB* colony.

The repulsion of the surfactant layer by TOB-treated *P. aeruginosa* suggests that PQS alone can deflect the surfactant layer. We reasoned that PQS in sufficient concentrations could deflect the surfactant layer from approaching tendrils, based on the observation that PQS repels swarm tendrils ((Bru et al., 2019) and Fig. S1B). Indeed, IRIS imaging showed that PQS deflects the surfactant layer, causing a simultaneous change in tendril direction (Fig. 6A and Movie S4 in Supplementary Materials). Dimethyl sulfoxide (DMSO), the solvent in which PQS was dissolved, did not deflect the surfactant layer (Fig. S6A in Supplementary Materials). Concave features were observed in the surfactant layer in the differential IRIS image, consistent with the surfactant deflection that was observed with PDMS and TOB-treated *P. aeruginosa* (Fig. 5C). We measured the magnitude of surfactant deflection by PQS through image analysis (see Methods section and Fig. S6B in Supplementary Materials) and found that the surfactant deflection area and tendril repulsion radius both increased with increasing concentrations of PQS (r = 0.95) (Fig. 6B). This finding supports the hypothesis that PQS repels tendrils physically by disrupting the surfactant layer flow. Together, these data suggest that tendril repulsion by stressed *P. aeruginosa* populations is caused by physicochemical interactions between PQS and the surfactant layer. These interactions in turn alter the surfactant flow and tendril direction around stressed *P. aeruginosa* populations. We note that while surfactant layer deflection has a significant role in tendril repulsion, the data does not rule out a role for sensing of PQS by the *P. aeruginosa* cells within the tendrils. It is possible that PQS sensing could affect the surfactant production or composition that could ultimately affect tendril direction.

**Figure 6.**
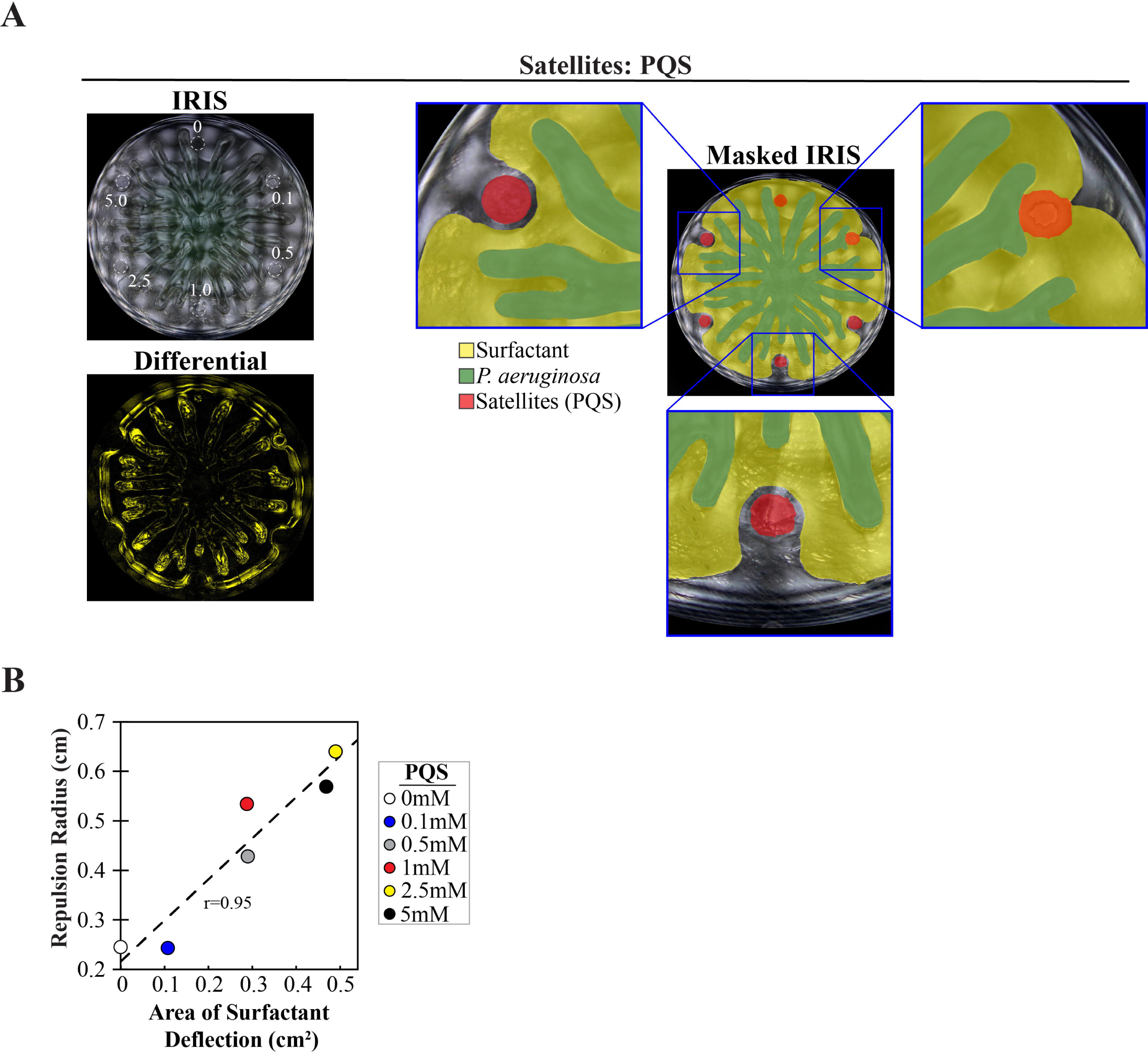
PQS increases both surfactant deflection and tendril repulsion. (A) IRIS, masked IRIS, and differential IRIS images of swarm interaction assays in which WT *P. aeruginosa* was spotted at the center and a range of PQS concentrations (0 - 5 mM) were spotted at satellite positions. Dashed lines in the IRIS image indicate the boundaries of the initial spots. Masked IRIS images indicate surfactant layer (yellow), *P. aeruginosa* (green), and initial PQS spots (red). The differential IRIS image indicates components of the surfactant layer and *P. aeruginosa* that are dynamic. Magnified IRIS images that are not masked are shown in Fig. S5B in the Supplementary Materials. (B) Tendril repulsion radii and surfactant deflection areas for the range of PQS concentrations tested. The correlation coefficient (r) value and a least squares fit (dashed line) are displayed on the plot. Repulsion radius was measured as the distance from the nearest tendril to the center of the PQS spot, regardless of whether the tendril contacted the boundary of the initial spot. Images were acquired 14.5 hours following inoculation.

### S. aureus repels P. aeruginosa tendrils by deflecting the surfactant layer

We rationalized that the repulsion of tendrils by *S. aureus* could be caused by a similar repulsion mechanism observed by PDMS, TOB-treated *P. aeruginosa*, and PQS. In particular, the amphipathic property of the PSM fibrils produced by *S. aureus* could deflect the *P. aeruginosa* surfactant layer, thereby altering tendril direction. IRIS imaging of *P. aeruginosa* swarms with *S. aureus* colonies at satellite positions revealed that the surfactant layer was indeed deflected and moved around WT *S. aureus* colonies (Fig. 7A-B, Fig. S5B and Movie S5 in Supplementary Materials). The *P. aeruginosa* tendrils moved within the boundary of the surfactant layer, resulting in the redirection of tendrils around the *S. aureus* colony. Surfactant deflection was significantly reduced in Δ*psmα* and Δ*psmα* Δ*hld* mutants (Fig. 7A-B), which are defective in the production of PSMs and do not repel tendrils. These data suggest that PSMs produced by *S. aureus* deflect the surfactant layer, which results in tendril repulsion.

**Figure 7.**
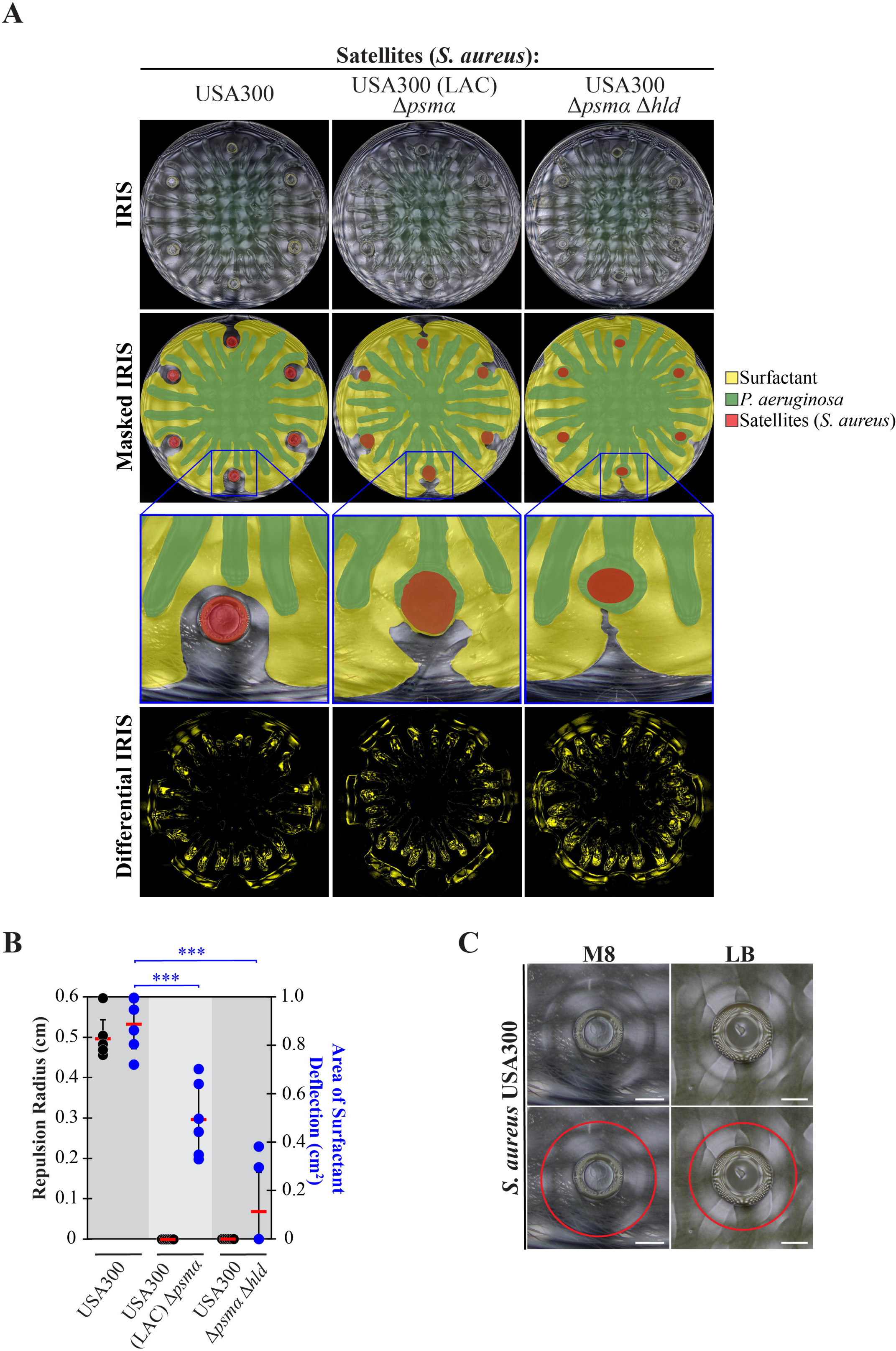
Surfactant layer deflection is diminished in PSM mutants. (A) IRIS, masked IRIS, and differential IRIS images of swarm interaction assays in which WT *P. aeruginosa* was spotted at the center and WT *S. aureus* (USA300) or PSM mutants were spotted at satellite positions. The masked IRIS images indicate the surfactant layer (yellow), *P. aeruginosa* (green), and initial spots (red). Differential IRIS images indicate components of the surfactant layer and *P. aeruginosa* that are dynamic. Magnified IRIS images that are not masked can be found in Fig. S5B in the Supplementary Materials. (B) Tendril repulsion radii (black dots) and surfactant deflection area (blue dots) of WT *S. aureus* and PSM mutants. The dots represent individual satellites, red bars indicate the average, and error bars indicate standard deviations. T-tests were performed as two-tailed distributions with unequal variance, *** denotes p < 0.001. (C) IRIS images of the fluidic layer produced by *S. aureus* USA300 cultured on M8 or LB medium containing 0.5% agar. The fluidic boundary is outlined in the lower images (red circle). Scale bar represents 3 mm. Images were acquired 14 hours following inoculation.

Additional support for this interpretation is the observation of a fluidic layer that develops outwards around a WT *S. aureus* colony during growth. The fluidic layer surrounds the colony and extends 3 mm beyond the colony edge (Fig. 7C and Movie S5 in Supplementary materials). While the full composition of the fluidic layer is unclear, we observed amyloid fibrils at a comparable distance (1.5 to 3 mm) from the *S. aureus* colony using TEM (Fig. 3B). The observation of the fluidic layer at a comparable distance as the fibrils offers a potential explanation for how the fibrils could be transported away from the colony edge. The fibrils are absent at this distance in the Δ*psmα* Δ*hld* mutant (Fig. 3B), which is consistent with the model that PSM fibrils mediate the deflection of the *P. aeruginosa* surfactant layer. We note that concave features in the surfactant layer were observed in the differential IRIS images near WT *S. aureus* colonies (Fig. 7A). The features are similar to the concave features produced by other tendril-repelling molecules (Fig. 5C and 6A). The size of the concave features is diminished in the Δ*psmα* Δ*hld* mutant compared to WT, though they are not entirely abolished (Fig. 7A). Together, these data suggest that PSM amyloid fibrils deflect the *P. aeruginosa* surfactant layer, resulting in the repulsion of *P. aeruginosa* tendrils. Because surfactant deflection is not completely abolished by the Δ*psmα* Δ*hld* mutant, this suggests that other molecules produced by *S. aureus,* including PSMβ could contribute to surfactant deflection. The deflection of surfactant by *S. aureus* is consistent with the mechanism of tendril repulsion observed by PDMS, TOB-treated *P. aeruginosa*, and PQS (Fig. 5C and 6A).

## Discussion

A fundamental aspect of multispecies bacterial communities is how one organism interacts with another. Here, we have investigated the spatial interaction between two competing opportunistic pathogens of major clinical importance, *P. aeruginosa* and *S. aureus*. Swarms of *P. aeruginosa* were repelled by *S. aureus*, which created a cell-free physical barrier and facilitated coexistence between the two species. We found that physicochemical interactions between the surfactant produced by *P. aeruginosa* and *S. aureus* PSMs create the separation. Our results provide insight into *P. aeruginosa* – *S. aureus* interactions and *P. aeruginosa* swarming and have important implications on the understanding of how bacteria interact with other species and host environments.

### Differential responses to antibiotic treatment

Our survey of clinical isolates revealed that under TOB-induced stress, *P. aeruginosa* and *S. aureus* have different effects on approaching *P. aeruginosa* swarms. Nearly all of the TOB-treated *P. aeruginosa* isolates repelled *P. aeruginosa* swarms (Fig. 1E-F). The converse was true in *S. aureus* strains. While almost all untreated clinical *S. aureus* strains repelled *P. aeruginosa* swarms, TOB treatment of these strains abolished the repulsive effect (Fig. 2D-E). This suggests that the antibiotic stress response of *P. aeruginosa* induces production of the signaling molecule PQS. However, an analogous stress response in which additional signaling molecules are produced was not observed in *S. aureus*.

### Repulsion of P. aeruginosa by S. aureus

The combination of IRIS and TEM imaging revealed a novel *P. aeruginosa - S. aureus* interaction. IRIS images showed that *S. aureus* colonies produce a fluidic layer that moves radially away from the colony, forming a ring up to 3 mm beyond the colony’s edge after 10 hours of growth. The fluidic layers are not visible under ambient lighting or with conventional imaging methods that use non-reflective illumination (Fig. 5B). TEM showed that PSM-dependent fibrils lie within the region containing the fluid, up to 3 mm away from the colony’s edge (Fig. 3B). The combination of IRIS and TEM data raises a number of questions regarding how fibrils are formed and are transported. It is possible that PSMs could be carried by the fluid and subsequently nucleate into fibrils during transport away from the colony, or that already-formed fibrils could be carried away by the fluidic layer. We point out that the fluidic layer produced by *S. aureus* is not limited to the growth conditions on swarm media plates here. In particular, a fluidic layer is produced by *S. aureus* that are cultured on LB plates (Fig. 7C), which are used widely for this organism.

We consider how PSMs deflect the *P. aeruginosa* surfactant layer. The surfactant consists of RLs and HAAs, which contain long chain hydrophobic domains. It would be expected that the surfactant layer of *P. aeruginosa* would merge with molecules of similar properties, such as the amphiphilic molecule PQS. In fact, RLs solubilize PQS (Calfee et al., 2005). Surprisingly, the surfactant layer is repelled by PQS (Fig. 6A). One possibility of how an amphiphilic molecule could repel the surfactant layer is that an intermediate layer of water provides a physical barrier between the surfactant layer and amphiphilic molecule. In the case of PSMs, which are amphipathic, a water layer could form between the surfactant and PSM fibrils, which would deflect the surfactant and produce a cell-free zone of repulsion between *S. aureus* and *P. aeruginosa* (Fig. 8A). An additional factor that could affect surfactant deflection is viscosity. We found that PDMS of lower viscosity (200 cSt) repelled tendrils to a greater extent than higher viscosity (1000 cSt) PDMS (p=0.02) (Fig. S4B in Supplementary Materials). Thus, a change in the viscosity of the area outside of the *S. aureus* colony produced by PSMs could contribute to surfactant layer deflection. In addition, small differentials in the amphipathicity between the surfactant layer and PSMs could also contribute to deflection. Finally, *S. aureus* may produce additional molecules that disrupt the surfactant layer that have not been identified in this study.

**Figure 8.**
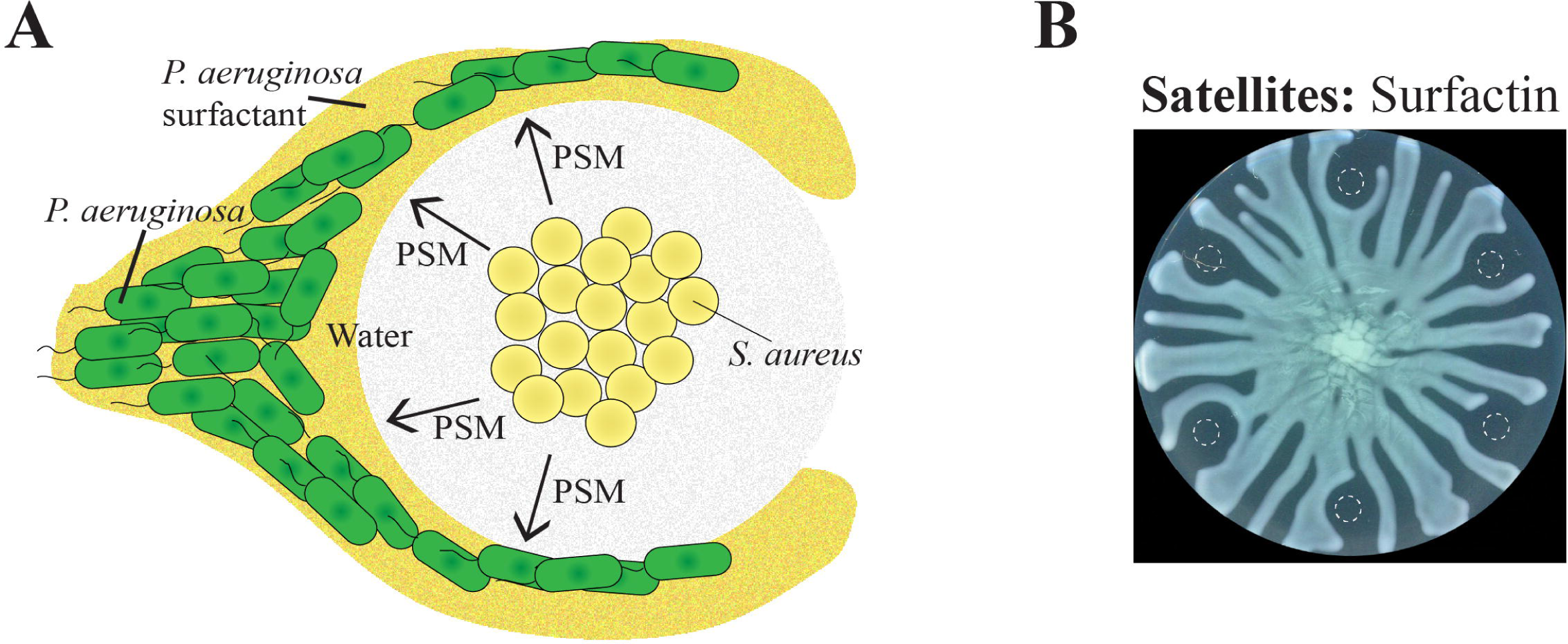
*P. aeruginosa* repulsion mediated by the surfactant layer. (A) Schematic of our *P. aeruginosa* swarm repulsion model. PSMs surround the *S. aureus* and deflect the *P. aeruginosa* surfactant layer, thereby causing tendril repulsion. (B) Swarming interaction assay in which WT *P. aeruginosa* and 10 µM surfactin naturally produced by *Bacillus subtilis* was spotted at the center and satellite positions, respectively. Images were acquired 16.5 hours following inoculation.

Previous investigations of *P. aeruginosa* - *S. aureus* interactions suggest that *P. aeruginosa* is predisposed to interact with *S. aureus* if the two species are within close proximity (microns) of one another (Limoli et al., 2019). During chronic infections, *P. aeruginosa* and *S. aureus* can evolve a mutualistic relationship in which *P. aeruginosa* does not inhibit *S. aureus* growth, but rather the interaction enhances *P. aeruginosa* growth (Michelsen et al., 2014; Frydenlund Michelsen et al., 2016). In contrast, our study shows that at the longer (millimeter) scale, *S. aureus* repels *P. aeruginosa* and the two species remain isolated, promoting species heterogeneity. In particular, RLs interact with the *S. aureus* membrane, making it more permeable to TOB, and thus sensitize *S. aureus* to TOB-mediated killing (Yarlagadda and Wright, 2019). The separation mechanism described here creates a physical barrier that reduces the likelihood that RLs reach *S. aureus* at the millimeter scale.

### Role of surfactant flow in swarm organization

Our analysis of *P. aeruginosa* swarming through IRIS imaging suggests that fluid mechanics have a significant role in tendril formation. In particular, the surfactant and tendrils form two separate layers that are coupled in motion, where the former is required for the movement of the latter. The surfactant layer constrains the movement of the swarm tendrils such that *P. aeruginosa* cannot move beyond the surfactant layer boundary. The movement of the surfactant layer around repulsive molecules such as PQS and PSMs thus directs the motion of the tendrils to move around these molecules. This mechanism has important implications for understanding how tendrils move on surfaces and are repelled or merge with other tendrils, such as is observed with the Δ*sadB* and Δ*rhlC* mutants (Caiazza et al., 2005). In particular, the flow of surfactant may need to be considered in such cases and in other aspects of swarm development.

While our results demonstrate an important role of the surfactant layer in tendril development, they do not preclude a role for biological sensing. Multiple *P. aeruginosa* sensors can explain the detection of molecules such as RLs and PQS by swarm tendrils. For example, it is possible that sensing of PSMs and PQS could alter the motion of tendrils around *S. aureus* and TOB-treated *P. aeruginosa*, respectively. However, a mechanism that explains how the tendril motion is altered in response to sensing is lacking.

In addition, we note that coupling between the surfactant and tendril layers is likely not the only mechanism that directs tendril movement. For example, after the surfactant layer reaches the boundary of the Petri dish, tendrils continue to move, although at a slower rate. Thus, decoupling between the surfactant and tendril layers is possible. Biophysical models of pressure-driven flow and Marangoni flow (Fauvart et al., 2012; Yang et al., 2017) could potentially explain the continued expansion of swarm tendrils in the absence of surfactant flow.

### Potential role of surfactant interactions in coexistence

In infection settings, *P. aeruginosa* and *S. aureus* frequently colonize the same environment. Here, it may be mutually beneficial for both species to remain spatially separated. *P. aeruginosa* may limit interactions with other organisms through surfactant production, while similarly, *S. aureus* may limit such interactions through PSM production, creating a barrier to surfactant flow. Consistent with this idea, significant levels of RLs have been measured in the lungs of CF patients (Kownatzki et al., 1987). Likewise, PSMs have a major role in lung infection models of *S. aureus* (Bloes et al., 2017). The repulsive interactions observed in this study could thus facilitate the coexistence of the two species in different micro-niches during coinfection. In particular, PSM fibrils could reduce RL-induced aminoglycoside sensitivity.

The finding that the *P. aeruginosa* surfactant layer has a major role in swarm organization may have implications on *P. aeruginosa* colonization in natural and host environments. The findings suggest that molecules which interact with *P. aeruginosa* surfactants can impact the spatial organization of the population. Moreover, the physicochemical nature of this interaction suggests that chemically-diverse hydrophobic molecules could alter this spatial organization. In support of this claim, we found that surfactin, a surfactant produced by *B. subtilis*, also repels *P. aeruginosa* swarms (Fig. 8B). This observation is remarkable because *B. subtilis* is evolutionarily distant from *P. aeruginosa*, yet causes repulsion of *P. aeruginosa* tendrils. In host environments, human lung cells produce a number of surfactants (Han and Mallampalli, 2015). In light of the results presented here, such surfactants could impact colonization or swarm-dependent spreading of *P. aeruginosa* during infection. In a clinical setting, patients suffering from genetic surfactant deficiency or premature newborns could be particularly sensitive to *P. aeruginosa* lung colonization if lung surfactants deflect *P. aeruginosa* rhamnolipids and swarming. Indeed, a mouse model of surfactant deficiency shows high susceptibility to *P. aeruginosa* colonization (Glasser et al., 2008). Additionally, given that numerous bacterial species found in natural and host environments produce unique surfactants, our findings raise the possibility that surfactants produced among different species of bacteria in polymicrobial environments could have a large impact on bacterial spatial organization, dissemination, and treatment outcomes.

## Materials & Methods

### Growth conditions and materials

Strains were streaked from frozen stocks onto LB Broth-Miller (BD, Franklin Lakes, NJ) plates containing 2% Bacto agar (BD) and incubated overnight at 37°C. Single colonies were inoculated into the same broth and incubated 16 – 18 hours in a roller drum at 20 rpm and 37°C. All strains used in this study are described in Table S1 in the Supplementary Materials. Clinical isolates were obtained from the Whiteson Lab and UCI Health Medical Microbiology Laboratory; no identifying information was collected and therefore IRB approval was not required.

The compounds in swarm interaction assays (Fig. 4); 200 cSt PDMS, 1000 cSt PDMS, oleic acid, linoleic acid, glyceryl trioleate, glyceryl trilinoleate, Triton X-100, Tween-20, and 2-heptyl-3-hydroxy-4-quinolone (PQS) were obtained from Sigma Aldrich (St. Louis, MO). Triton X-100 and Tween-20 were used at 0.2% and 2%, respectively, due their inhibitory growth effect on *P. aeruginosa* swarms at higher concentrations. PQS was dissolved in dimethylsulfoxide (DMSO). Strains at satellite positions were treated with antibiotics using tobramycin (Thermo Fisher Scientific Hampton, NH) at a concentration of 0.5 mg/mL.

### Swarm interaction assays

Assays were performed as described previously (Bru et al., 2019, 2020). Briefly, 100 x 15mm Petri dishes contained 20 mL of M8 minimum medium (Bru et al., 2020) supplemented with 1 mM MgSO_4_, 0.2% glucose, 0.5% Casamino Acids (BD), and 0.5% Bacto agar (BD). Following sterilization, media in Petri dishes solidified for 1 hour with lids on at room temperature, and then dried for 30 minutes without lids in a laminar flow hood at 300 cubic ft. / min. with approximately 40 to 50% ambient humidity. Strains were cultured 16 – 18 hours in LB broth. WT *P. aeruginosa* was spotted at the center of the swarming plate in a 5 µL droplet. Strains or compounds were spotted at satellite positions using 6 µL droplets at 6 satellite positions that were equidistant from each other along a 5.8 cm-diameter circle centered at the swarming plate (Fig. S6C in the Supplementary Materials). Plates were incubated overnight at 37°C with a darkened Petri dish lid. Images were acquired every 30 minutes for 18 – 20 hours using an Epson scanner (Epson, Los Alamitos, CA) that was controlled using RoboTask (robotask.com) and then processed using ImageJ 1.54d (NIH, Bethesda, MD). For the antibiotic treatment of strains at satellite positions, tobramycin (Thermo Fisher Scientific) was mixed with the bacterial inoculum to a final concentration of 0.5 mg/mL and the mixture was spotted immediately onto swarming plates in 6 µL droplets. The repulsion radius at each satellite colony was measured as the distance from the center of the satellite to the nearest tendril along a line that is tangent to the 5.8 cm-diameter circle that is concentric with the swarming plate (Fig. S6C in the Supplementary Materials). If a tendril contacted the boundary of the initial satellite spot, the repulsion radius was marked as zero, except for the measurements to determine the correlation between repulsion radius and surfactant deflection area by PQS (Fig. 6B).

### Transmission electron microscopy

Mesh copper grids that were 3 mm in diameter (Electron Microscopy Sciences, Hatfield, PA) were placed with the carbon layer facing up at or adjacent to inoculation positions on swarming plates. 6 μL of *S. aureus* that had been cultured in LB broth for 16 – 18 hours was spotted onto the inoculum position. Plates containing the inoculums and copper grids were incubated at 37°C and 50% humidity for 16 - 18 hours. The grids were stained using 1% uranyl acetate (Electron Microscopy Sciences), dried overnight at room temperature, and imaged using a JEM-2800 High Throughput Electron Microscope (JEOL, Akishima, Japan) using an acceleration voltage of 200 kV.

### Contact angle measurements

Test compounds were pipetted vertically in one microliter droplets at room temperature onto a Notak oleophobic-coated (SilcoTek, Bellefonte, PA) polished stainless steel sheet surface and imaged immediately for 10 seconds using a Model 90 goniometer (ramé-hart, Succasunna, NJ) with a Summit SK2-3.1X 3.1MP digital camera (OptixCam, Roanoke, VA). Contact angles were measured from timelapses at the 7 second timepoint after the initial droplet placement onto the surface using ImageJ and the sessile drop method. Angles were measured for the left and right sides of the droplet in the images and averaged.

### IRIS imaging

Imaging was performed as described previously (Kasallis et al., 2023). Briefly, swarming plate surfaces were illuminated by an ASUS LCD VE278Q monitor (ASUS, Fremont, CA) that displayed a structured image that consisted of evenly-spaced white squares in a 17 x 31 grid pattern on a black background. The reflection of the structured image on the swarming plate surface was acquired using a Canon EOS Rebel T7 DSLR (Canon, Melville, NY) with a Canon EF-S 18-55 mm lens. The inner surfaces of Petri dishes were scratched with a metal wire brush to reduce reflections. Swarming plates were incubated at 37°C and 50% humidity. For timelapses, the lid was removed and replaced at regular intervals using a robotic arm to allow for image acquisition. Masks of the IRIS images were constructed using the magnetic lasso tool in Adobe Photoshop (Adobe, San Jose, California). Differential images were constructed using the imabsdiff function in Matlab version R2017a (Mathworks, Natick, MA) using images that were acquired at 5 minute (Fig. 5B) or 30 minute intervals (all others). The surfactant deflection area was defined as the total area in the vicinity of the satellite spot that did not contain *P. aeruginosa* surfactant (Fig. S6B in the Supplementary Materials). Our identification of the *P. aeruginosa* surfactant layer boundary was aided by following surfactant flow in timelapse IRIS movies and using differential IRIS images. An additional boundary of the deflection area was defined as an arc that connected two of the nearest surfactant layer boundaries that were not deflected.

## Author contributions

AS, JLB, and QZ developed the IRIS device; JLB, SK, KW, NMHK, DHL, and AS designed experiments and determined project direction; JLB, SK, RC, QZ, PP, and JN performed experiments and analyzed data; JLB, SK, and AS drafted and edited the manuscript; all authors reviewed the manuscript.

## Acknowledgements

We thank Joshua Shrout for helpful discussions about swarming and Cassiana Bittencourt (UCI Health) and Zachary Lu, Joann Phan and Tara Gallagher (Whiteson Lab) for collecting the clinical isolates. N.M.H-K. was supported by a grant from the Independent Research Fund Denmark (1054-00099B) and A.S. was supported by a grant from the NIH NIAID (R56AI163196).

## Notes

### Competing Interest Statement

The authors have declared no competing interest.

