## Supplementary Materials for "The great divide: rhamnolipids mediate separation between *P. aeruginosa* and *S. aureus*"

Supplementary Movie Legends
Supplementary Figures S1-S6
Supplementary Table S1
Supplementary References

**Supplementary Movie Legends**

**Movie S1. *P. aeruginosa* swarm interaction assays using1000 cSt PDMS at satellite positions.** IRIS (left) and differential IRIS (right) timelapses of swarm interaction assays in which *P. aeruginosa* and 1000 cSt PDMS were spotted at the center and satellite positions, respectively. Individual images were captured every 30 minutes following inoculation over the course of 18 hours.

**Movie S2. *P. aeruginosa* swarm interactions assays using tobramycin-treated *P. aeruginosa* at satellite positions.** IRIS (left) and differential IRIS (right) timelapses of swarm interaction assays in which *P. aeruginosa* was spotted at the center and *P. aeruginosa* that were mixed with tobramycin to a concentration of 0.5 mg/mL and spotted as 6 µL droplets at satellite positions. Images were captured every 30 minutes following inoculation over the course of 18 hours.

**Movie S3. *P. aeruginosa* swarm interaction assays using *P. aeruginosa* Δ*rhlAB* at satellite positions.** IRIS (left) and differential IRIS (right) timelapses of swarm interaction assays in which wild-type *P. aeruginosa* and the *P. aeruginosa* Δ*rhlAB* mutant were spotted at the center and satellite positions, respectively. Images were captured every 30 minutes following inoculation over the course of 18 hours.

**Movie S4. *P. aeruginosa* swarm interaction assays using PQS at satellite positions.** IRIS (left) and differential IRIS (right) timelapses of swarm interaction assays in which wild-type *P. aeruginosa* was spotted at the center and PQS (0 - 5 mM) was spotted at the satellite positions. Images were captured every 30 min following inoculation over the course of 18 hours.

**Movie S5. *P. aeruginosa* swarm interaction assays using *S. aureus* at satellite positions.** IRIS (left) and differential IRIS (right) timelapses of swarm interaction assays in which wild-type *P. aeruginosa* and wild-type *S. aureus* strain USA300 were spotted at the center and satellite positions, respectively. Images were captured every 30 minutes following inoculation over the course of 18 hours.

**Supplementary Figures S1-S6**

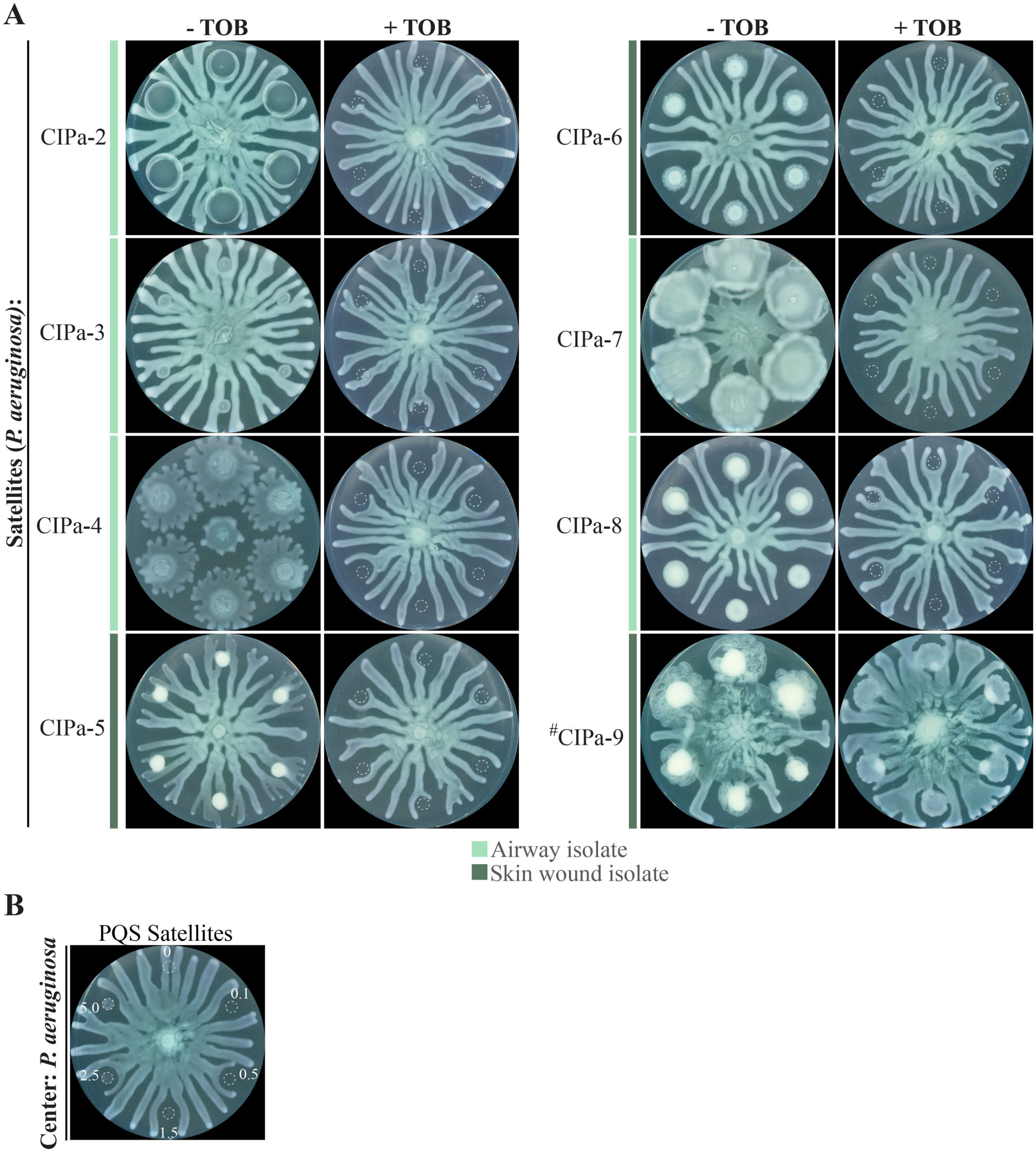


**Figure S1. Swarm repulsion by clinical isolates of *P. aeruginosa*.** (A) Swarm interaction assays in which wild-type *P. aeruginosa* and clinical isolates of *P. aeruginosa* (CIPa) were spotted at the center and satellite positions, respectively. CIPa strains were spotted with or without tobramycin (TOB). Tobramycin (TOB) treatment was performed by mixing TOB with bacteria to a final concentration of 0.5 mg/mL and spotting 6 μL of the mixture onto the swarm plate. Color bars indicate if the strains were isolated from the airway or skin wound. *CIPa-9 was resistant to TOB. (B) Swarm interaction assay in which wild-type P. aeruginosa was spotted at the center and PQS (0 – 5 mM) was spotted at satellite positions. Dashed lines indicate the boundaries of initial inoculum spots. Images were acquired 18-20 hours following inoculation.

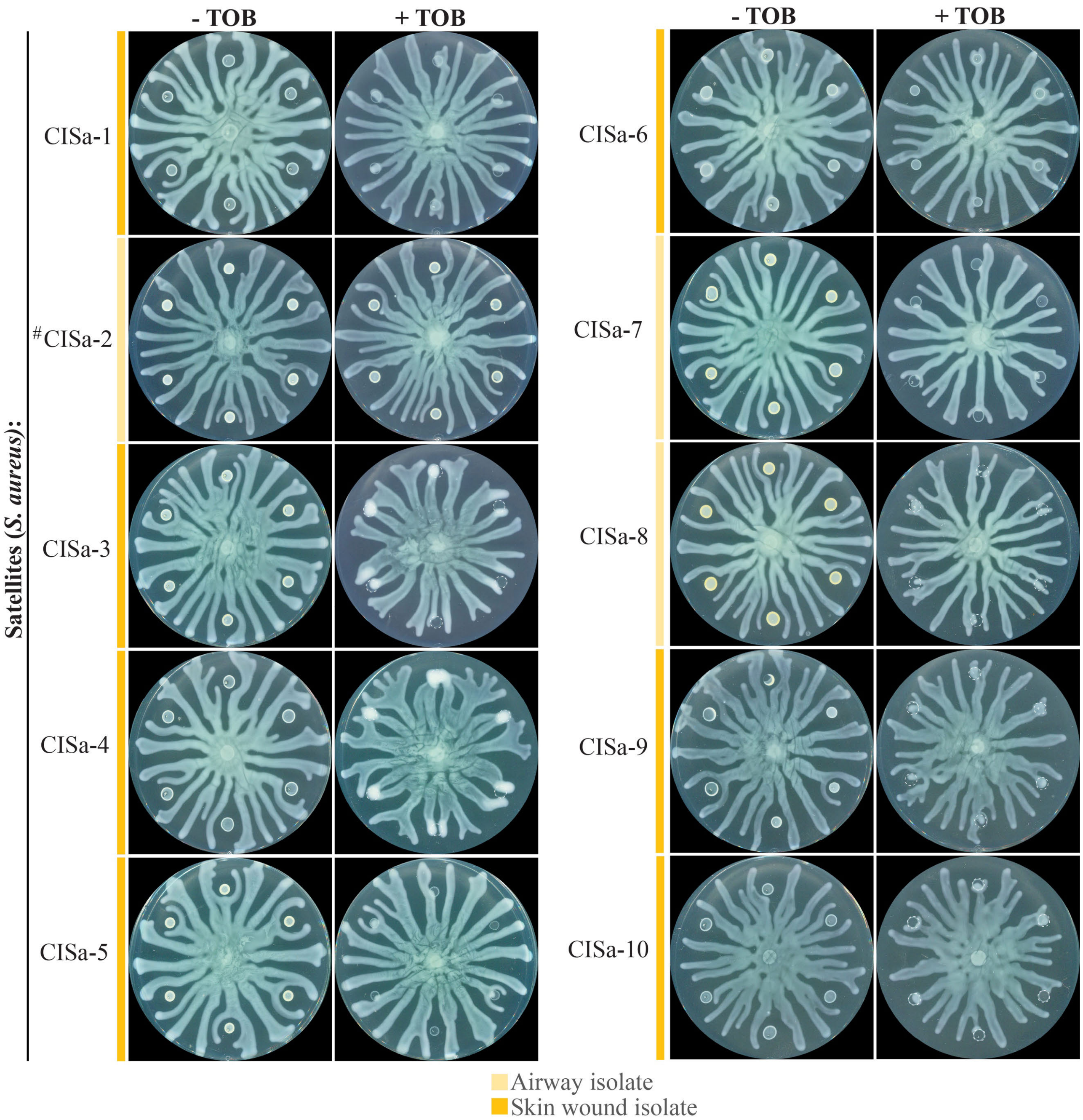


**Figure S2. Swarm repulsion by clinical isolates of *S. aureus*.** Swarm interaction assays in which wild-type *P. aeruginosa* and clinical isolates of *S. aureus* (CISa) were spotted at the center and satellite positions, respectively. CISa strains were spotted with or without tobramycin (TOB). Tobramycin (TOB) treatment was performed by mixing TOB with bacteria to a final concentration of 0.5 mg/mL and spotting 6 µL of the mixture onto the swarm plate. Color bars indicate if the strains were isolated from the airway or skin wound. *CISa-2 was resistant to TOB. Images were acquired 18-20 hours following inoculation.

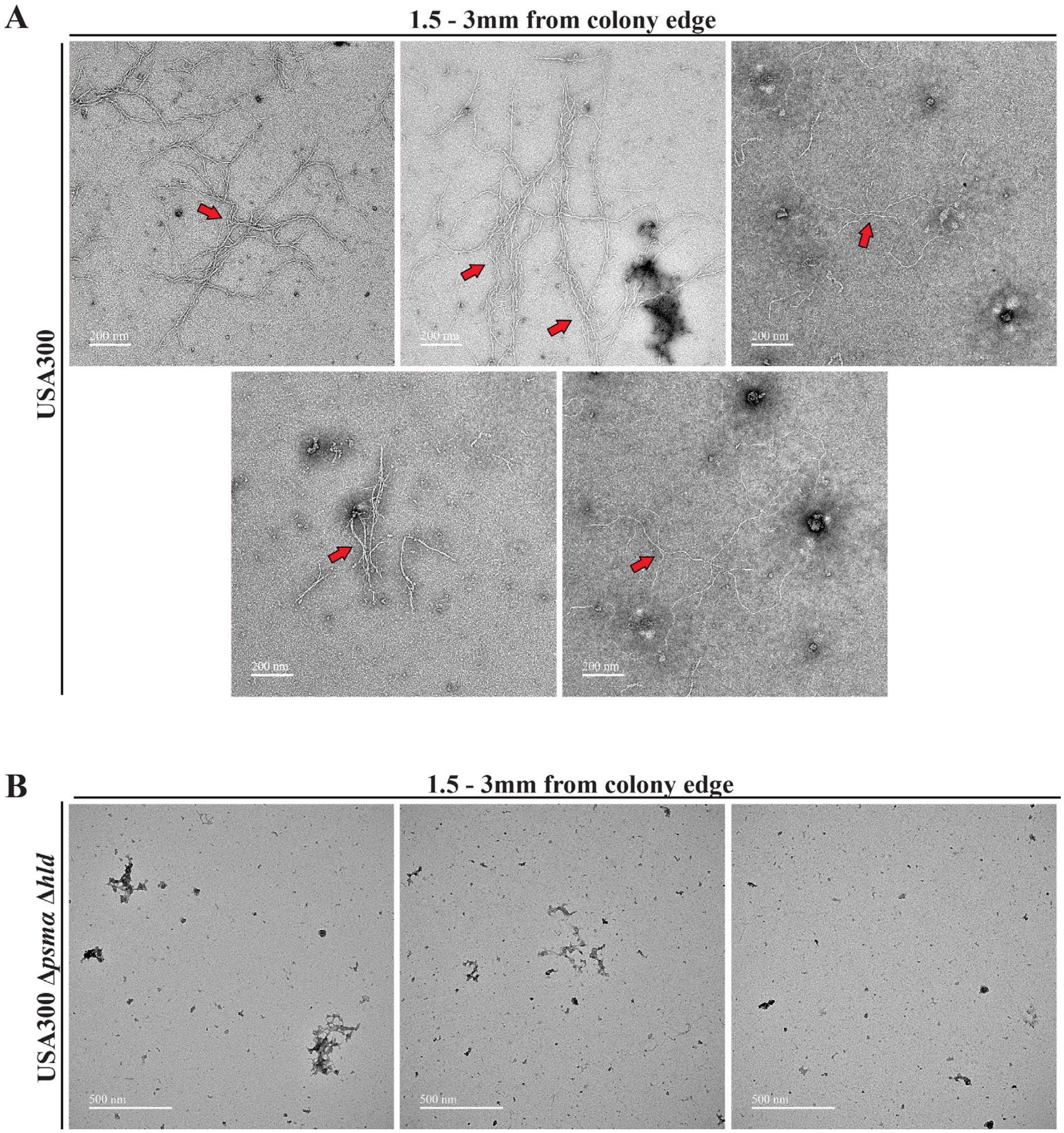


**Figure S3. Transmission electron microscopy (TEM) images in vicinity of *S. aureus* colonies.** TEM images of areas that were between 1.5 and 3mm from the edge of colonies of (A) wild-type *S. aureus* strain USA300 and (B) the *S. aureus* Δ*psmα* Δ*hld* mutant. Scale bars indicate 200 nm in (A) and 500 nm in (B).

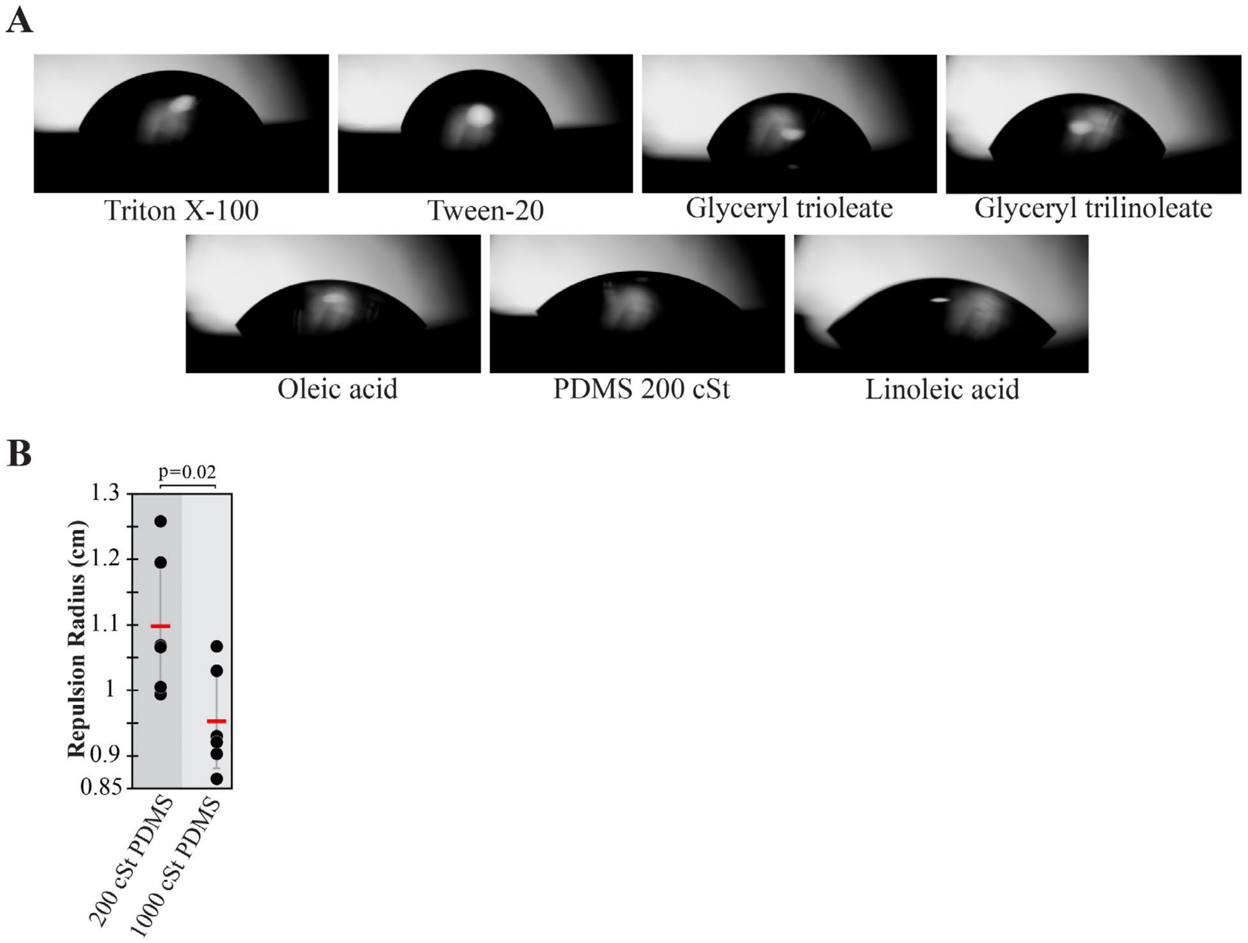


**Figure S4. Contact angles and repulsion radii of hydrophobic molecules.** (A) Images of hydrophobic molecules on an oleophobic surface acquired by a contact angle goniometer. Triton X-100 and Tween-20 were used at concentrations of 0.2% and 2%, respectively. (B) Tendril repulsion radii by 200 cSt and 1000 cSt viscosities of PDMS. Measurements were performed on images that were acquired 15 hours following inoculation. Red lines indicate average repulsion radius and error bars indicate standard deviation. T-tests were performed as two-tailed distributions with unequal variance.

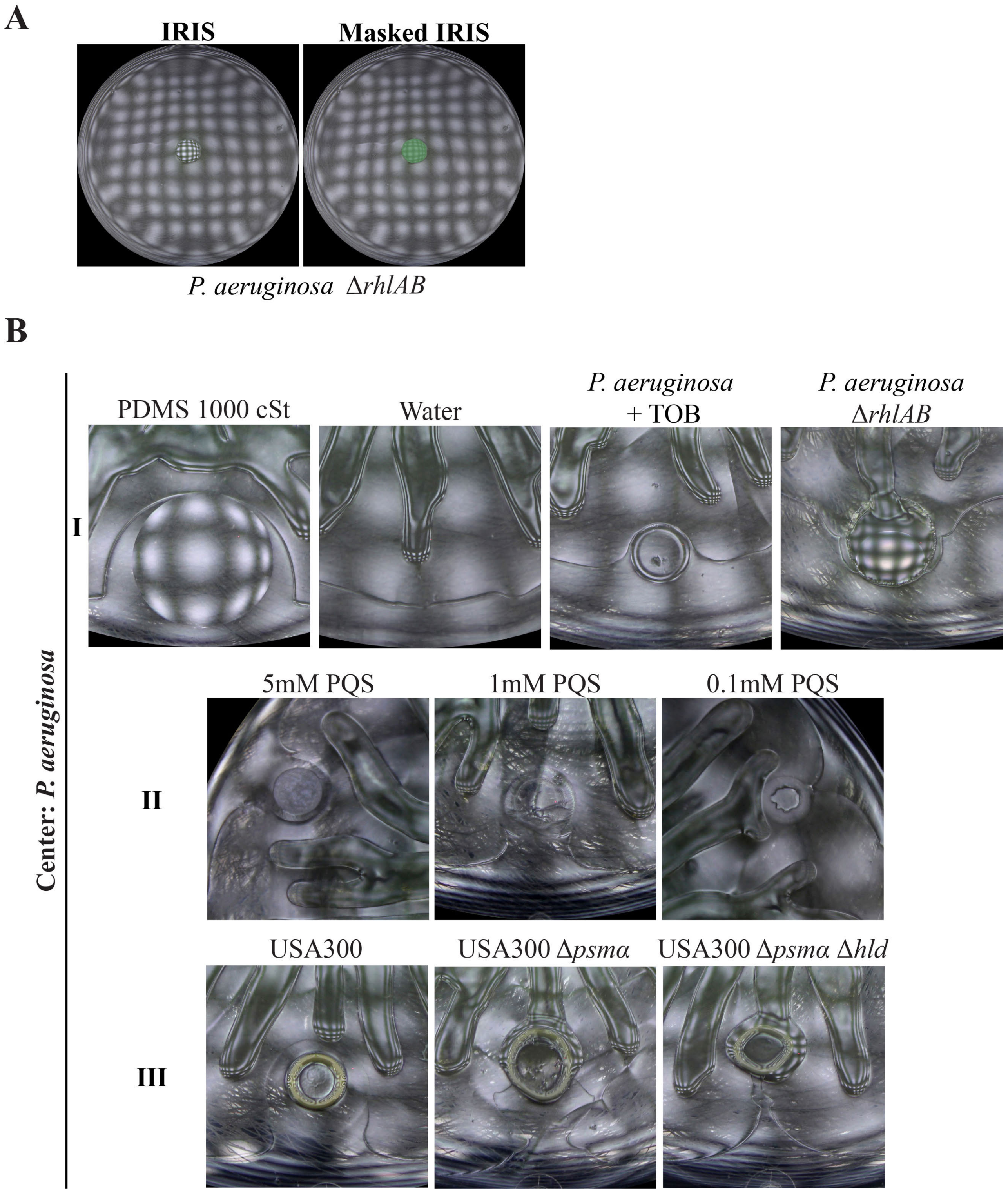


**Figure S5. Surfactant production and surfactant interactions.** (A) IRIS image (left) and masked IRIS image (right) of *P. aeruginosa* Δ*rhlAB* (green) spotted at the center of the plate and imaged after 20 hours of growth. A surfactant layer is not detected. (B) IRIS images in which wild-type *P. aeruginosa* is spotted at the center and test strains or compounds are spotted at satellite positions. The images in I, II, and III are used to construct the masked IRIS images in Figs. 5C, 6A, and 7A, respectively.

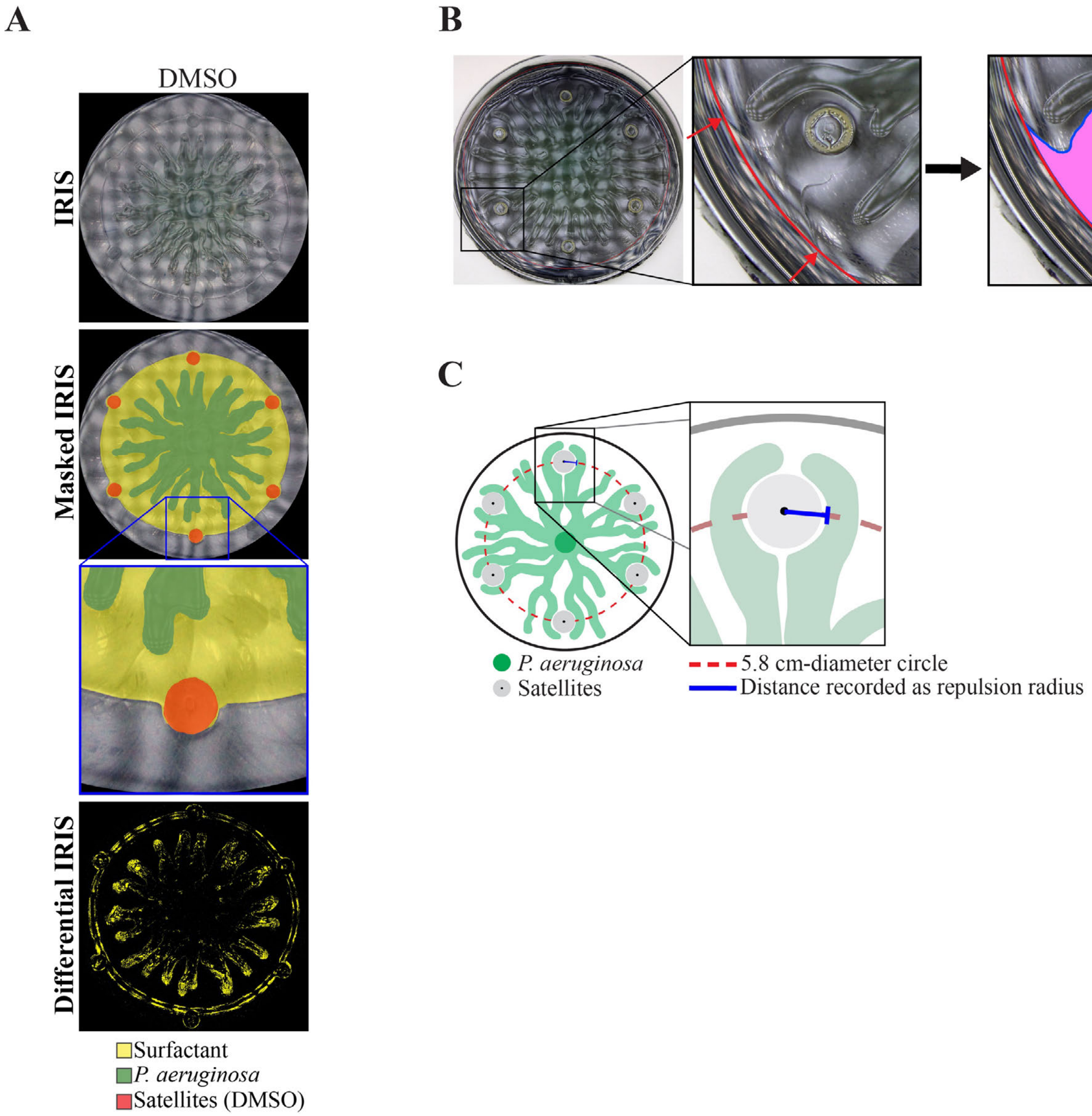


**Figure S6. Swarm interactions with DMSO and surfactant and tendril measurement methods.** (A) IRIS, masked IRIS, and differential IRIS images of *P. aeruginosa* and DMSO that are spotted at the center and at satellite positions, respectively. Images were acquired 14 hours following inoculation. Masked IRIS images indicate the surfactant layer (yellow), *P. aeruginosa* (green), and the initial boundaries of the satellite spots (red). (B) Graphic depicting how the surfactant deflection area was determined, using *S. aureus* USA300 at the satellite position as an example. The boundary of the surfactant layer near the satellite position was identified (blue line). An arc (red) that connects two of the nearest surfactant layer boundaries that are not deflected was defined as an additional boundary. The surfactant deflection area (pink) was defined as the area enclosed by these boundaries. (C) Schematic depicting how tendril repulsion radius is determined. Test strains or compounds (gray) were spotted along a 5.8 cm-diameter circle (red dashed line) that is concentric with the swarming plate. The repulsion radius (blue line) at each satellite colony was measured as the distance from the center of the satellite position (black dot) to the nearest tendril along a line that is tangent to the circle. If a tendril contacted the boundary of the initial satellite spot, the repulsion radius was recorded as zero.

### Supplementary Table S1 – Strains used in this study

| Name | Lab Strain Name | Strain | Description | References | Source |
|---|---|---|---|---|---|
| CIPa-1 | PAnmFLR01 (P1) | *P. aeruginosa* | Airway isolate | (Quinn et al., 2016) | Whiteson lab |
| CIPa-2 | PAmFLR02 (P2m) | *P. aeruginosa* | Airway isolate | (Quinn et al., 2016) | Whiteson lab |
| CIPa-3 | WI 1-2 | *P. aeruginosa* | Airway isolate | This study | UCI Health |
| CIPa-4 | WI 4-7 | *P. aeruginosa* | Airway isolate | This study | UCI Health |
| CIPa-5 | WI 12-22 | *P. aeruginosa* | Skin wound isolate | This study | UCI Health |
| CIPa-6 | WI 14-26 | *P. aeruginosa* | Skin wound isolate | This study | UCI Health |
| CIPa-7 | WI 16-30 | *P. aeruginosa* | Airway isolate | This study | UCI Health |
| CIPa-8 | WI 17-32 | *P. aeruginosa* | Airway isolate | This study | UCI Health |
| CIPa-9 | WI 20-38 | *P. aeruginosa* | Skin wound isolate | This study | UCI Health |
| CISa-1 | WI 2-4 | *S. aureus* | Skin wound isolate | This study | UCI Health |
| CISa-2 | WI 6-10 | *S. aureus* | Airway isolate | This study | UCI Health |
| CISa-3 | WI 7-12 | *S. aureus* | Skin wound isolate | This study | UCI Health |
| CISa-4 | WI 9-15 | *S. aureus* | Skin wound isolate | This study | UCI Health |
| CISa-5 | WI 10-18 | *S. aureus* | Skin wound isolate | This study | UCI Health |
| CISa-6 | WI 12-21 | *S. aureus* | Skin wound isolate | This study | UCI Health |
| CISa-7 | WI 13-24 | *S. aureus* | Airway isolate | This study | UCI Health |
| CISa-8 | WI 15-28 | *S. aureus* | Airway isolate | This study | UCI Health |
| CISa-9 | WI 19-36 | *S. aureus* | Skin wound isolate | This study | UCI Health |
| CISa-10 | WI 21-40 | *S. aureus* | Skin wound isolate | This study | UCI Health |
| USA300 | SADHL129 | *S. aureus* | USA300 | (Chaney et al., 2017) | Wozniak lab |
| USA300 Δ*psmα* Δ*hld* | SADHL130 | *S. aureus* | USA300 Δ*psmα*1-4 *δ*ATG – ATT | (Syed et al., 2015) | Wozniak lab |
| USA300 (LAC) | SADHL121 | *S. aureus* | USA300 (LAC) | (Centers for Disease Control and Prevention (CDC), 2003; McDougal et al., 2003) | Alex Horsewill lab |
| USA300 (LAC) Δ*psmα* Δ*hld* | SADHL137 | *S. aureus* | USA300 (LAC) Δ*psmα*1-4 *δ*ATG – ATT | | Alex Horsewill lab |
| USA300 (LAC) Δ*psmα* | SADHL138 | *S. aureus* | USA300 (LAC) Δ*psmα*1-4 | | Alex Horsewill lab |
| USA300 (JE2) | SADHL43 | *S. aureus* | USA300 (JE2) derivative | (Fey et al., 2013) | Cheung lab |
| Wild-type *P. aeruginosa* | AFS27E.1 | *P. aeruginosa* | PA14 *att*Tn7::[P$_{A1/04/03}$-*mCherry*] *aacC1*::FRT | (Bru et al., 2019) | Siryaporn lab |
| *P. aeruginosa* D*rhlAB* | BR04.1 | *P. aeruginosa* | PA14 Δ(*rhlAB*)::FRT | (Bru et al., 2019) | Siryaporn lab |
| *P. aeruginosa* D*rhlAB* D*pqsA* | AFS82.1 | *P. aeruginosa* | PA14 Δ(*rhlAB*)::FRT Δ(*pqsA*)::FRT | (Bru et al., 2019) | Siryaporn lab |
